# BMP-dependent patterning of ectoderm tissue material properties modulates lateral mesendoderm cell migration during early zebrafish gastrulation

**DOI:** 10.1101/2023.07.21.550024

**Authors:** Stefania Tavano, David B. Brückner, Saren Tasciyan, Xin Tong, Roland Kardos, Alexandra Schauer, Robert Hauschild, Carl-Philipp Heisenberg

**Affiliations:** Institute of Science and Technology Austria, Klosterneuburg, Austria; Center for Pathobiochemistry and Genetics, Medical University of Vienna, Vienna, Austria

## Abstract

Cell migration is a fundamental process during embryonic development. Most studies on cell migration *in vivo* have focussed on the migration of cells using the extracellular matrix (ECM) as their substrate for migration. In contrast, much less is known about how cells migrate on other cells, as found in early embryos when the ECM has not yet formed. Here, we show that lateral mesendoderm (LME) cells in the early zebrafish gastrula use the overlying ectoderm as their substrate for migration. We show that the lateral ectoderm is permissive for the animal pole-directed migration of LME cells, while the ectoderm at the animal pole of the gastrula halts animal pole-directed LME migration. These differences in the permissive properties of the ectoderm for LME migration are due to the lateral ectoderm being more cohesive and viscous than the animal ectoderm. Consistently, tuning cell contractility in lateral and animal ectoderm via modulation of the actomyosin cytoskeleton is sufficient to change their permissiveness for LME migration. Finally, we found that BMP signalling is critical for reducing animal ectoderm cohesion and, thus, its capacity to halt LME migration towards the animal pole. Collectively, these findings identify a critical role of ectoderm tissue cohesion and viscosity in guiding LME migration during zebrafish gastrulation.

## Introduction

Cell migration is a fundamental process in organism development and homeostasis, and defects in its regulation are at the basis of various diseases. The extracellular environment plays a pivotal role in the regulation of cell migration. Work *in vitro* and *in vivo* has shown that exposure to pre-existing or self-generated gradients of biochemical signals can guide cells over long distances^1, 2^. Moreover, physical confinement can influence the migratory behaviour of cells and direct their migration^3, 4^. An essential component of the extracellular environment is the extracellular matrix (ECM), on which cells can migrate. Especially the biochemical composition and mechanical properties of the ECM, i.e., its adhesiveness, stiffness and topology, have been shown to affect cell migration both *in vivo* and *in vitro*^5, 6^. In contrast, much less is known about how cells migrate without ECM.

An *in vivo* model of ECM-independent cell migration is lateral mesendoderm (LME) migration in the early zebrafish gastrula. In the developing zebrafish embryo, the onset of ECM deposition coincides with the start of gastrulation^7, 8^. However, until late gastrulation, ECM does not form a distinct and coherent network^7, 8^. Thus, LME cell migration in the early zebrafish gastrula occurs largely without ECM. LME migration is characterised by three distinct phases (Figure 1A): upon ingression at the germ ring margin, LME progenitors collectively migrate towards the animal pole of the gastrula as a loosely connected population between the yolk cell membrane and the overlying ectoderm ^9,10^. For moving towards the animal pole, LME cells use a self-generated gradient of *toddler*^11^ (mesoderm) and random walk^10, 12^ (endoderm), with these two modes of cell migration being coordinated by Cxcl12b/Cxcr4a signalling^13, 14^. After a period of persistent motion, the animal movement of LME cells ceases at mid-gastrulation, when LME cells enter into a second phase characterised by uncoordinated tumbling movements and little displacement^15, 16^. This phase is followed by a third phase of convergence movements, where LME cells undergo highly persistent and directed migration towards the forming body axis^9^. This third migration phase depends on Bone Morphogenetic Protein (BMP)^17^ and non-canonical Wnt-signalling^18^ and is guided by a gradient of Cadherin-mediated cell-cell adhesion^19^ and requires the interaction with ECM^8^.

**Figure 1.**
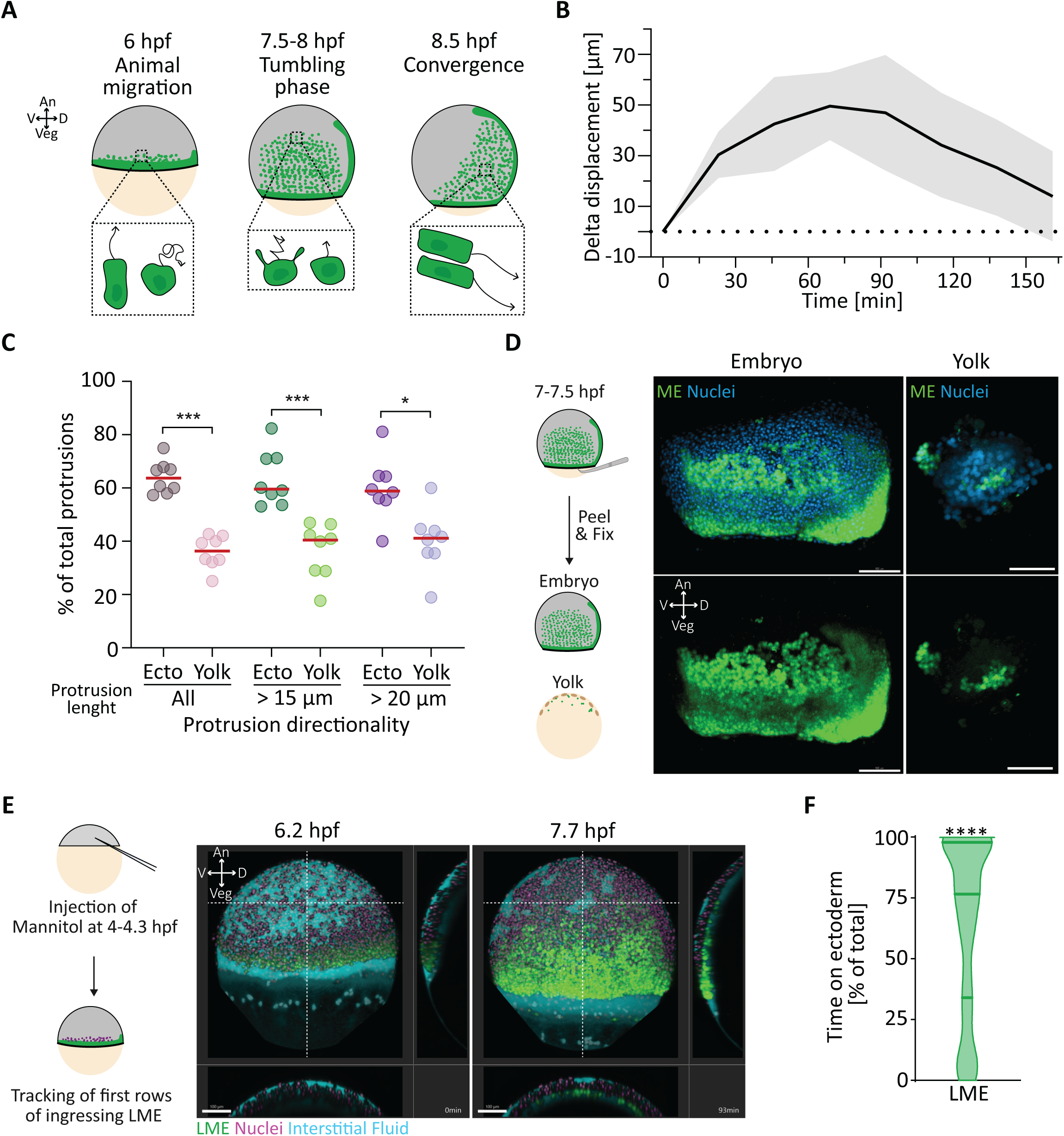
Ectoderm is the primary substrate for lateral mesendoderm animal pole-directed migration. (A) Schematic representation of lateral mesendoderm (LME) migration during zebrafish gastrulation. An, animal; Veg, vegetal; D, dorsal; V, ventral; hpf, hours post-fertilization. (B) Delta displacement of the LME front over time. The displacement of the LME front from the origin of migration towards the animal pole was tracked every 23 min (t0 ∼6.2 hpf) for at least 115 min. The delta displacement was computed as the difference of each time point from the previous one (Y-axis). Values are shown as mean (solid black line) with standard deviation (SD, light grey area). Dotted line represents 0 μm on the Y-axis. Number of embryos, 11. See also Figure S1 and Video S1. (C) Directionality of LME actin-positive protrusions during animal pole-directed migration. LME cells were sparsely labelled by injection of 7.5 pg *LifeAct-RFP* mRNA into a single blastomere at the 64/128 cells stage (2-2.25 hpf). Protrusion directionality was quantified for an average of 50 min of migration (∼2 min 14 sec frame rate, t0 ∼6.2 hpf). Scatter dot plot shows the mean of each cell during migration. Red line represents the median. Protrusion length is quantified from the cell centre. Number of cells, 8; number of embryos, 2. Total number of protrusions: all, 2778 (14.1 ± 1.6 SD protrusions per cell per frame); total length from the cell centre > 10 μm, 2392 (12 ± 0.9 SD protrusions per cell per frame); total length the cell centre > 20 μm, 437 (2.2± 1.1 SD protrusions per cell per frame). Statistical test, Mann- Whitney test; ***p<0.001, *p<0.05. See also Video S2. (D) Yolk peeling assay. On the left, schematic representation of the yolk peeling assay. The yolk membrane was manually peeled from the blastoderm of *Tg(sebox::eGFP)* embryos at 65- 70% epiboly (7-7.5 hpf) using forceps and both embryo and yolk were immediately fixed in 4% PFA. Blastoderm and yolk were then stained with DAPI (nuclei) before imaging. On the right, representative image of blastoderm and yolk. Green, eGFP (mesendoderm, ME); cyan, DAPI (nuclei). Scale bar, 100 μm. An, animal; Veg, vegetal; D, dorsal; V, ventral. (E) Mannitol-induced detachment of the blastoderm from the yolk. On the left, schematic representation of the experimental setup. Sphere/dome stage (4-4.3 hpf) *Tg(sebox::eGFP); Tg(actb2::H2A-mCherry)* embryos were injected with 1 nl of 350 mM Mannitol together with 1.25 µg/µl 10.000 MW Alexa Fluor™ 647 dextran to label the interstitial fluid. First ingressing LME cells (∼6.2 hpf) (for definition see Figure S1B and Methods) were tracked for ∼92 min from the onset of animal pole-directed migration (∼2 min frame rate). On the right, still images of LME cells (lateral view) from a representative timelapse movie at the start (left, 6.2 hpf) and end (right, 7.7 hpf) of the migration period. White dashed lines indicate the position of the yz (right) and xz (bottom) cross-sections. Green, eGFP (LME cells); magenta, H2A- mCherry (nuclei); cyan, Alexa Fluor™ 647 dextran (interstitial fluid). Scale bar, 100 μm. An, animal; Veg, vegetal; D, dorsal; V, ventral. See also Video S3. (F) Time LME cells use the ectoderm as substrate for migration. LME cells were scored as either on the ectoderm or on the yolk membrane at each timepoint for ∼92 min from the onset of animal pole-directed migration (6.2 hpf). Data are shown as percentage of the total time of migration. Number of cells, 163; number of embryos, 3. Statistical test, One sample Wilcoxon test; ****p<0.0001. See also Figure S2.

While past work has provided insight into the molecular and cellular regulation of both animal pole-directed and convergence LME cell migration, comparably little is yet known about why LME cells cease moving towards the animal pole at mid-gastrulation and, instead, enter into a tumbling phase followed by convergence movements. Given that ECM at mid- gastrulation does not yet show any distinct accumulation within the embryo^7, 8^, changes in the interaction between LME cells and their surrounding tissues, i.e. the underlying yolk cell and overlying ectoderm germ layer, are likely to be involved in this process. Here, we show that LME preferentially uses the ectoderm as a substrate for their animal pole-directed migration and that BMP-dependent patterning of ectoderm material properties restricts animal pole- directed migration of LME cells.

## Results

### Ectoderm is the primary substrate for lateral mesendoderm migration

To understand why LME cells stop migrating towards the animal pole and, instead, turn dorsally, we first analysed how cells at the leading edge of the LME migrate towards the animal pole (6.2 hours post fertilisation (hpf) - 8.8 hpf, Figures 1B and S1A, Video S1) in *tg(Sebox::eGFP)* embryos, a fish line expressing cytosolic GFP under the control of a pan- mesendodermal marker gene promoter^20^. Consistent with previous observations^11, 15, 16^, we found cells at the LME front first move towards the animal pole (Figures S1A, 0 – 92 min), as evidenced by an initial constant increase of their displacement (Figure 1B, 0 – 69 min), and then abruptly stop migrating and transit into a tumbling phase at mid-gastrulation (Figure S1A, 115 - 161 min), recognisable by a sharp decrease in their displacement (Figure 1B, 115 – 161 min). This confirms the observation that the ability of LME cells to move animally sharply decreases at mid-gastrulation^15^. To further understand the migratory behaviour of LME cells before the tumbling phase (6.2-7.7 hpf), we analysed their migration at the single-cell level. Interestingly, we found that the first LME cells reaching the animal part of the embryo at 7.7 hpf (“most animal”, Figure S1B) display migratory properties different from cells following behind (more vegetally) (Figure S1B), as evidenced by a higher persistence and animal-ward directionality of their migration and, consequently, overall higher displacement along the animal-vegetal (AnVeg) axis (Figure S1C-F, LME). To determine whether this behaviour also applies to lateral endoderm cells, which as part of the LME cells have been previously shown to move via random motion^12^, we analysed their migratory behaviour during the first phase of animal migration using a fish line expressing cytosolic GFP under the control of an endoderm marker gene promoter^21^. We found that lateral endoderm cells behave similarly to all LME cells (Figure S1B-F, Endo), with the first cells reaching the animal part of the embryo displaying a more persistent and directed migration.

Previous studies have shown that differences in substrate properties, such as stiffness, can affect cell migration^1–3, 5, 6, 22^. To determine whether changes in substrate properties along the AnVeg axis might be responsible for the abrupt slowing-down of LME animal migration, we first investigated which substrate LME cells use for their migration. In zebrafish, ECM forms a coherent network between the forming germ layers only at the end of gastrulation^7, 8^ and, thus, is unlikely to serve as a substrate for LME migration at the onset of gastrulation. Without ECM, LME cells can only choose between two possible substrates, the overlaying ectoderm and the underlying yolk cell (Figure S1G). To assess which of these possible substrates are used by LME cells, we first analysed the preferred orientation of actin protrusions of LME cells during their animal migration (Figure 1C and Video S2), focusing on actin-rich protrusions that were clearly recognizable and the length of which was more the three-times the average nuclear radius (> 15 µm). Interestingly, most protrusions were preferentially oriented towards the ectoderm, suggesting that LME cells are in contact with the overlying ectoderm rather than the underlying yolk cell (Figure 1C). To corroborate these findings, we sought to probe whether LME cells preferentially adhere to the ectoderm or the yolk cell by mechanically separating the yolk cell from the blastoderm at 65-70% epiboly, followed by immediate fixation of both (Figure 1D). After separation, we observed that LME cells were found mainly on the ectoderm rather than the yolk cell, suggesting that they preferentially adhere to the ectoderm. To further challenge this suggestion, we tested which substrate LME cells prefer if unable to touch both substrates simultaneously. To this end, we injected 350 mM Mannitol (an inert sugar) between ectoderm cells at the sphere-to-dome stage (4-4.3 hpf), which we previously showed to increase interstitial fluid accumulation at the yolk cell-to-ectoderm boundary^23^ (Figure 1E and Video S3). LME cells migrating into this enlarged fluid-filled space had to choose between the ectoderm and yolk cell as their migration substrate but could not simultaneously adhere to both substrates (Figure 1E and Video S3). In Mannitol-injected embryos, LME cells displayed animal migration indistinguishable from LME cells in uninjected control embryos (Figure S2A-D, LME). When analysing whether LME cells use yolk cell or ectoderm for their animal migration, we found that they preferred ectoderm (Figure 1F, median 76.1), suggesting that the latter is their primary substrate. Lateral endoderm cells, in contrast, could be subdivided into two roughly equal subpopulations (Figure S2E, median 54.3), one predominately using the ectoderm and the other the yolk membrane as their substrate for migration. Moreover, we found that lateral endoderm migration persistence and displacement along the AnVeg axis, but not their overall directionality, were reduced in Mannitol-injected embryos (Figure S2A-D, Endo), indicating that lateral endoderm cells require the close spatial proximity with both ectoderm and yolk cell for proper migration.

Finally, we asked whether and how migrating on the ectoderm affects LME and lateral endoderm animal migration. We found a positive correlation between displacement on the AnVeg axis and time spent migrating on the ectoderm for both LME cells, and also lateral endodermal cells as part of the LME (Figure 2A). To further corroborate these data, we separately analysed cells predominantly migrating on the ectoderm or the yolk cell (Figure 2B and S3A). Notably, we observed that LME and lateral endoderm cells had the highest displacement and persistence when migrating on the ectoderm (Figures 2C-D and S3C-D). Moreover, cells using mainly the ectoderm showed a higher animal directionality and displacement along the AnVeg axis than cells using the yolk cell as their migrational substrate (Figures 2E-F and S3E-F). Collectively, this suggests that LME cells preferentially use the overlying ectoderm as their substrate for migration, and that migrating on the ectoderm promotes LME and lateral endoderm animal pole-directed migration.

**Figure 2.**
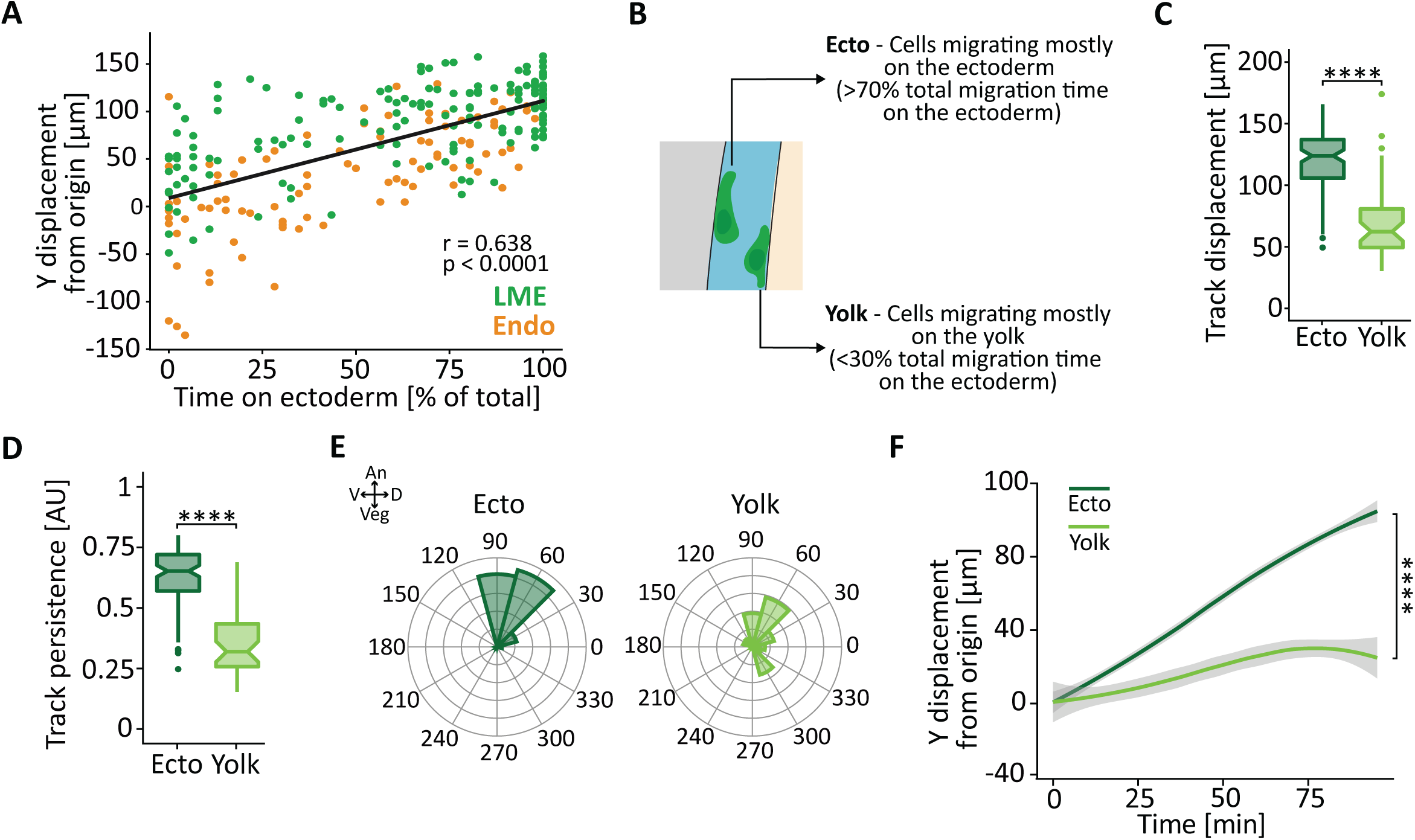
Lateral mesendoderm cells migrating on the ectoderm display most pronounced animal pole-directed migration. (**A-F**) Sphere/dome stage (4-4.3 hours post fertilization, hpf) *Tg(sebox::eGFP); Tg(actb2::H2A- mCherry)* embryos (lateral mesendoderm (LME), A-F) or *Tg(sox17::eGFP)* embryos injected with 50 pg *H2A-mCherry* mRNA (nuclei) at the 1-cell stage (lateral endoderm (Endo), A) were injected with 1 nl of 350 mM Mannitol together with 1.25 µg/µl 10.000 MW Alexa Fluor™ 647 dextran to label the interstitial fluid. First ingressing cells (for definition see Figure S1B and Methods) were tracked for ∼92 min from the onset of animal pole-directed migration (∼2 min frame rate, t0 ∼6.2 hpf). See also Figure S2 and S3. (A) Correlation between the time LME and Endo cells migrate on the ectoderm (X-axis) and their displacement along the animal-vegetal (AnVeg) axis, expressed as final Y-displacement from the origin (Y-axis). Green dots, LME; orange dots, Endo cells. Statistical test, Spearman r. Number of pairs, 259 (163 LME cells from 3 embryos; 96 Endo cells from 2 embryos). See also Figure S2. (B) Explanation of the subgrouping used for the analyses in (C-F) and in Figure S3. Cells using ectoderm as substrate for either more than 70% or less than 30% of their migration time were defined in the analysis as cells migrating mostly on the ectoderm (Ecto) or mostly on the yolk cell (Yolk), respectively. See also Figure S3. (C) Total displacement of LME cells using mostly the ectoderm (Ecto, dark green) or the yolk cell (Yolk, light green) as substrate for their migration. Number of cells: Ecto, 99; yolk, 42. Number of embryos, 3. Statistical test, Mann-Whitney test: ****p<0.0001. (D) Migration persistence of LME using mostly the ectoderm (Ecto, dark green) or the yolk cell (Yolk, light green) as substrate for their migration. Number of cells: Ecto, 99; yolk, 42. Number of embryos, 3. Statistical test, Mann-Whitney test: ****p<0.0001. (E) Migration directionality of LME cells using mostly the ectoderm (Ecto, dark green) or the yolk cell (Yolk, light green). Directionality is shown as rose diagram where each concentric circle represents a different frequency range, with 0 at the centre and increasing by 10 for each subsequent circle. An, animal; Veg, vegetal; D, dorsal; V, ventral. Number of cells: Ecto, 99; yolk, 42. Number of embryos, 3. (F) Displacement of LME cells towards the animal pole over time. The displacement along the AnVeg axis is quantified as cumulative Y-displacement from the origin of migration (Y-axis) and displayed as smooth plot. Solid line represents the mean, grey ribbon displays confidence interval. LME mostly using the ectoderm (Ecto) or the yolk cell (Yolk) as substrate for their migration are shown in dark green and light green, respectively. Statistical test on the final Y displacement, Mann-Whitney test: ****p<0.0001.

### Ectoderm patterning along the AnVeg axis modulates LME animal pole-directed migration

Given the critical role of the ectoderm in promoting animal pole-directed migration of LME, we asked whether local differences in ectoderm properties along the gastrula AnVeg axis might be responsible for the sharp slowing-down of LME animal migration at mid-gastrulation (Figures 1B and S1A, Video S1). To address this possibility, we performed ectoderm tissue transplantation experiments, replacing lateral ectoderm with either animal (heterotypic transplantation) or lateral (homotypic transplantation) ectoderm (Figures 3A-B, S4A and Videos S4-5). As expected, homotypic transplantation of lateral-to-lateral ectoderm did not affect LME migratory behaviour compared to untransplanted embryos (Figure 3C-E). In contrast, heterotypic animal-to-lateral ectoderm transplants substantially decreased LME persistence and displacement along the AnVeg axis (Figure 3C-E), resulting in a substantial reduction in the ability of LME cells to reach the animal part of the gastrula before entering into the tumbling phase (Figures 3F and S4A). This suggests that animal ectoderm is a less permissive substrate for LME animal pole-directed migration than lateral ectoderm.

**Figure 3.**
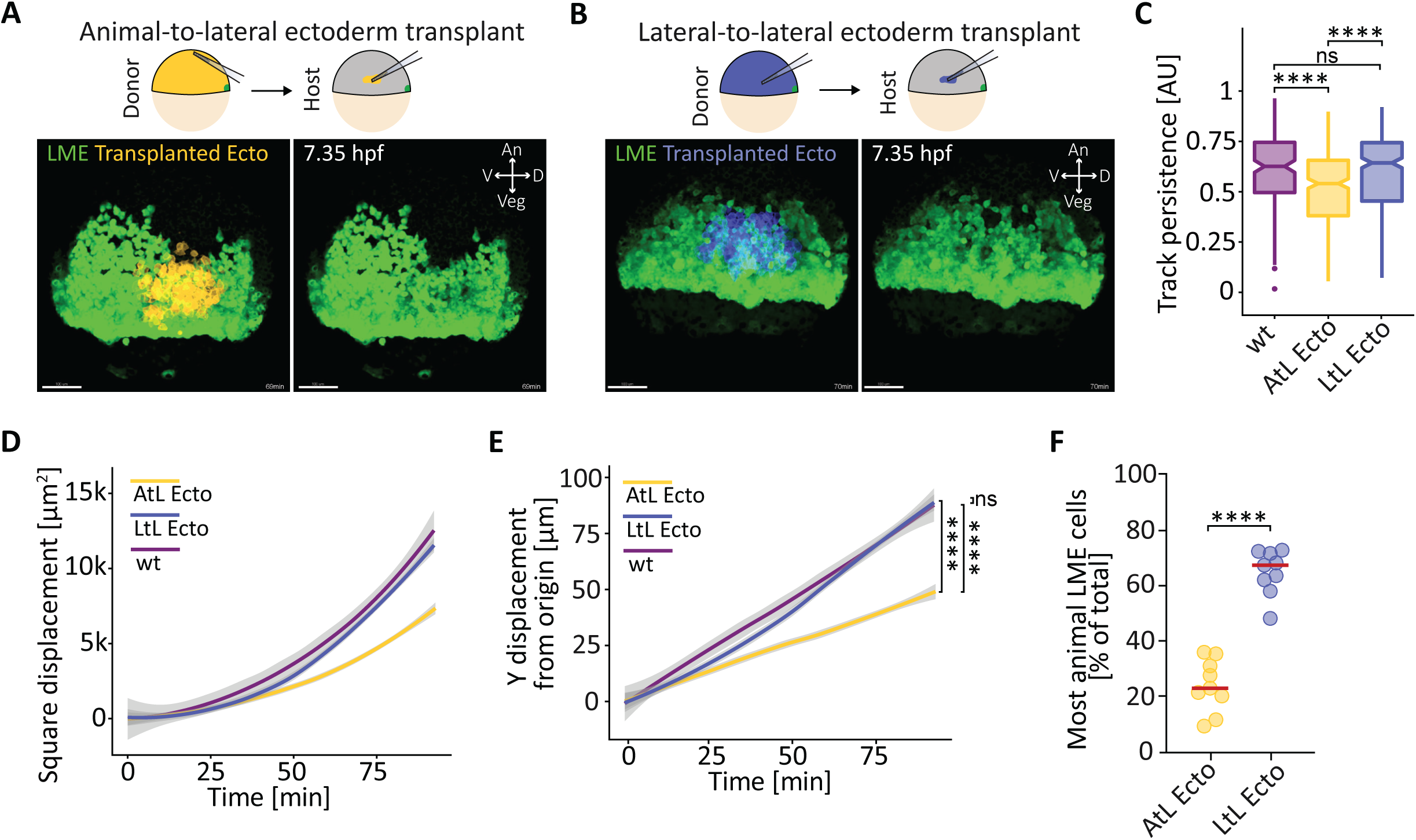
Animal ectoderm inhibits lateral mesendoderm animal pole-directed migration. (**A,B**) Animal-to-lateral (homotypic) and lateral-to-lateral (heterotypic) ectoderm transplants assay. Animal ectoderm (yellow, A) or lateral ectoderm (blue, B) from a wildtype *Tg(sebox::eGFP)* donor embryo injected with 10-15 pg *H2B-mKO2k* and 100 pg *membrane RFP* mRNAs to mark nuclei and plasma membrane, respectively, were transplanted at germ ring stage (5.7 hours post fertilization, hpf) in the lateral ectoderm of a stage-matched wildtype *Tg(sebox::eGFP)* host embryo (injected with 10-15 pg *H2B-mKO2k* mRNA (nuclei)). After transplantation, lateral mesendoderm (LME) cells encountering the transplanted ectoderm (for definition, see Figure S4 and Methods) were tracked for ∼92 min (∼2 min 11 sec frame rate, t0 ∼6.2 hpf). Upper part, schematic representation of the experimental setup. Lower part, fluorescence images of LME cells and transplanted animal (yellow, A) or lateral (blue, B) ectoderm (lateral view) from representative timelapses ∼70 min after the start of LME migration (∼7.35 hpf). Green, eGFP (LME cells); yellow, membrane RFP (transplanted animal ectoderm); blue, membrane RFP (transplanted lateral ectoderm). Scale bar, 100 μm. An, animal; Veg, vegetal; D, dorsal; V, ventral. See also Video S4 and S5. (**C**) Migration persistence of LME cells in transplanted versus non-transplanted wildtype (wt) embryos. Number of cells, 173 from 3 non-transplanted embryos (wt; purple), 327 from 9 embryos with animal-to-lateral ectoderm transplants (AtL Ecto, yellow), 327 from 9 embryos with lateral-to-lateral ectoderm transplants (LtL Ecto, blue). Dataset for wt corresponds to data shown in Figure S1. Statistical test, Mann-Whitney test: ns not significant, **** p<0.0001. See also Figure S4. (**D**) Square Displacement of LME cells in transplanted versus non-transplanted (wt) embryos displayed as smooth plot. Solid line represents the mean, grey ribbon displays confidence interval. Number of cells, 173 from 3 non-transplanted embryos (wt, purple), 327 from 9 embryos with animal-to-lateral ectoderm transplants (AtL Ecto, yellow), 327 from 9 embryos with lateral-to-lateral ectoderm transplants (LtL Ecto, blue). Dataset for wt corresponds to data shown in Figure S1. See also Figure S4. (**E**) Displacement of LME cells towards the animal pole over time cells in transplanted versus non-transplanted (wt) embryos. The displacement along the animal-vegetal (AnVeg) axis is quantified as cumulative Y-displacement from the origin of migration (Y-axis) and displayed as smooth plot. Solid line represents the mean, grey ribbon displays confidence interval. Number of cells, 173 from 3 non-transplanted embryos (wt, purple), 327 from 9 embryos with animal-to-lateral ectoderm transplants (AtL Ecto, yellow), 327 from 9 embryos with lateral- to-lateral ectoderm transplants (LtL Ecto, blue). Dataset for wt corresponds to data shown in Figure S1. Statistical test on the final Y-displacement, Mann-Whitney test: ns not significant, **** p<0.0001. See also Figure S4. (**F**) Percentage of most animally-migrating LME cells in transplanted embryos. Tracked LME cells were scored as most animally-migrating at the end of the tracked time (∼92 min, 7.7 hpf) if they were within a 100 µm-wide bin (along the AnVeg axis) from the most animally-located LME cells that had not encountered the transplanted ectoderm (see also Figure S4). Scatter dot plot shows the mean of each embryo. Red line represents the median. Number of embryos, 9 embryos with animal-to-lateral ectoderm transplants (AtL Ecto, yellow) and 9 with lateral-to-lateral ectoderm transplants (LtL Ecto, blue). Dataset for wt corresponds to data shown in Figure S1. Statistical test, Mann-Whitney test: **** p<0.0001.

Given that substrate material properties are thought to play a pivotal role in modulating cell migration^1, 3, 24^, we asked whether animal and lateral ectoderm might show different tissue material properties during gastrulation. To obtain insight into ectoderm tissue material properties, we first analysed how ectoderm thickness changes when spreading over the yolk cell during epiboly when animal pole-directed LME migration occurs (6.2 – 7.7 hpf, Figure 4A- C), a phenomenon we had previously associated with differences in blastoderm tissue rigidity in the early zebrafish embryo^25^. Interestingly, we found that the epibolizing ectoderm thins faster at the animal pole than near its margin (Figure 4B-C). To relate these findings to LME migration, we subdivided the ectoderm into ‘lateral’ ectoderm, being in contact with migrating LME, and ‘animal’ ectoderm, being devoid of migrating LME, and quantified their changes in tissue thickness during the period of LME animal migration. At the end of LME animal migration (7.7 hpf), the lateral ectoderm showed a decrease in thickness of 27.5% (0.735 ± 12.2 SD), while the animal ectoderm reduced their thickness by 62.7% (0.373 ± 7.5 SD) (Figure 4A-D). Together these data indicate that similar to the blastoderm at doming^25^, epibolizing ectoderm undergoes differential thinning.

**Figure 4.**
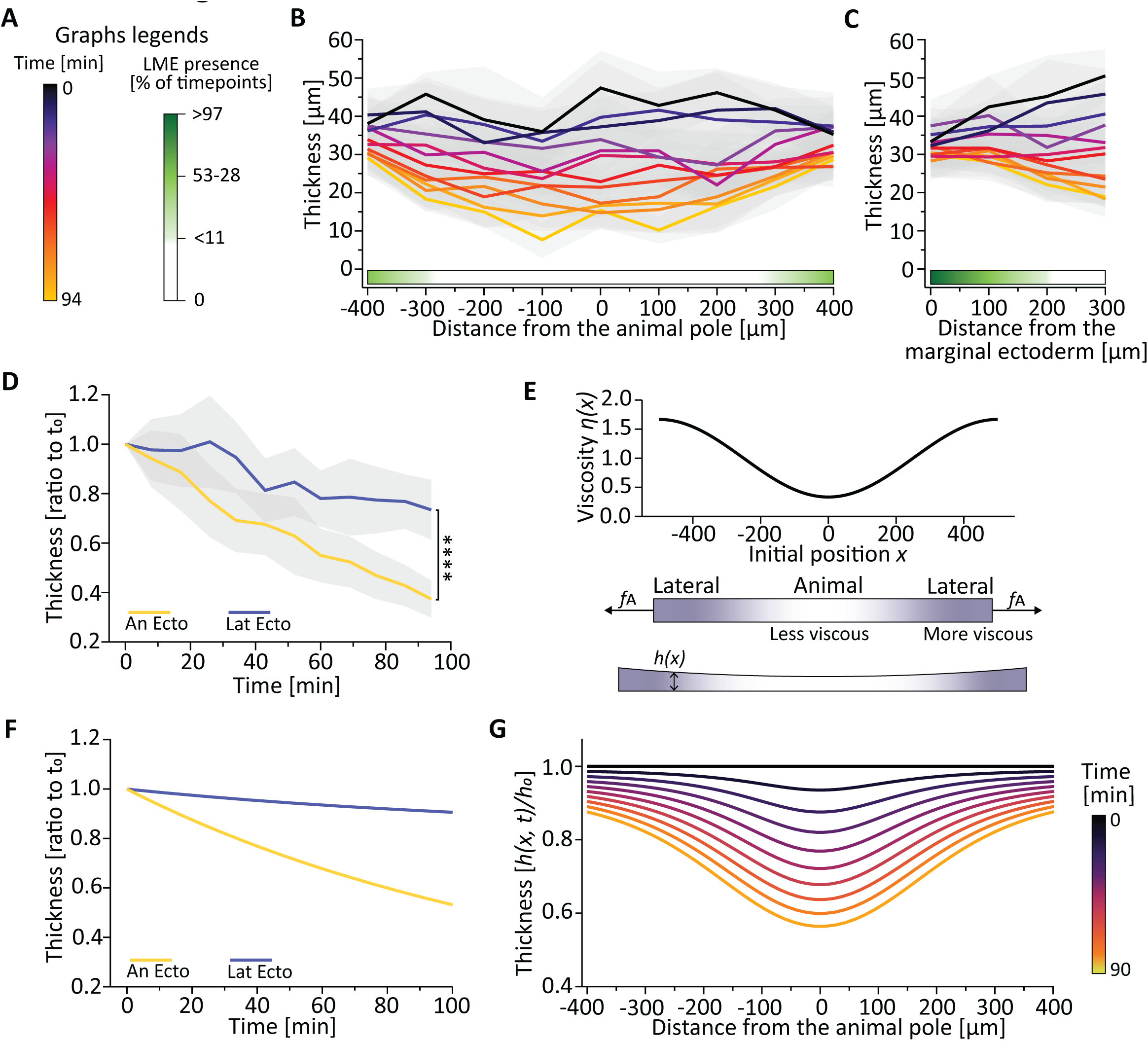
Lateral and animal ectoderm undergo differential thinning. (**A-D**) Ectoderm thickness as a function of developmental time in *Tg(sebox::eGFP);Tg(actb2:: mKate2-tpm3)* embryos injected with 1 nl of 1 µg/µl 10.000 MW Alexa Fluor™ 647 dextran to mark interstitial fluid and imaged from either the animal pole (B) or the lateral side (C) of the gastrula. The fluorescent signals of LME (eGFP), actin cytoskeleton (mKate2-tpm3) and interstitial fluid (Alexa Fluor™ 647 dextran) were used to create binary mask of the embryo, the LME and the interstitial fluid, which were then used to quantify the ectoderm thickness (y-axis) for 94 min from the onset of LME migration (6.2 – 7.7 hours post fertilization, hpf, fire colormap (A)). See also Video S6 and S7. (**B,C**) Ectoderm thickness at the animal pole and in 100 µm-wide bins lateral to the animal pole (x-axis) (B), and in the most marginal ectoderm (100-150 µm from the embryo equator) and in 100 µm-wide bins animal to the margin (x-axis) (C). Values are shown as mean (solid line) with standard deviation (SD, light grey area). Green gradient bar represents the presence of LME in a specific position. Number of embryos: animal pole view (B), 6; lateral view (C), 6. (**D**) Ectoderm thickness at the animal pole and in 100 µm-wide bins lateral to the animal pole and in at the most marginal ectoderm (100-150 µm from the embryo equator) and in 100 µm- wide bins animal to the margin. For each embryo, bins were defined as either lateral ectoderm (Lat Ecto, blue), in case the ectoderm was in contact with the LME at least for one timepoint during the imaging period, or animal ectoderm (An Ecto, yellow), in case the ectoderm was not in contact with LME during the entire imaging period. Thickness is shown as ratio to the thickness at the onset of LME migration (t0). Values are shown as mean (solid line) with standard deviation (SD, light grey area). Number of embryos, 12. Statistical test, Two-way ANOVA: **** p<0.0001. (**E-G**) Model of the ectoderm thickness as a function of developmental time. Ectoderm tissue is modelled as a passive fluid with patterned viscosity from the animal pole (lowest) to the margin (highest) and subjected to an external constant pulling force (*fA*), which cause an increase of tissue length with a speed of ∼1 µm/min (corresponding to the experimentally measured blastoderm epiboly speed) (E). In (F), animal (An Ecto, yellow) and lateral (Lat Ecto, blue) ectoderm thickness is shown as ratio to the thickness at t0. Animal ectoderm corresponds to 200 µm at the animal pole, while lateral ectoderm corresponds to the 200 µm at the margin. In (G), ectoderm thickness was quantified as a function of time (90 min, fire colormap) at the animal pole and 400 µm lateral to it (x-axis). See also Figure S4.

During epiboly, the thinning of the ectoderm occurs as an effect of the forces emanating from the underlying yolk syncytial layer (YSL), pulling the blastoderm margin towards the vegetal pole of the gastrula^26^. Those pulling forces are, therefore, expected to be larger near their origin (the margin) than away from it (the animal pole). Thus, the more pronounced tissue thinning at the animal pole is likely due to differential ectoderm tissue material properties rather than pulling forces along its AnVeg axis. To further conceptualize these findings, we developed a minimal ectoderm thinning model for investigating whether this differential ectoderm thinning could result from spatial differences in tissue viscosity^25^ (Figures 4E and S4B-G). We assumed that the ectoderm is a passive fluid with a conserved volume and that with epiboly progressing, the ectodermal tissue is subjected to an external pulling force causing its elongation with a constant speed of approximately 1.5 µm/min^27^. We first assumed equal viscosity throughout the ectoderm and asked how the tissue thickness changes over time. We found that the tissue thickness decreases equally in animal and lateral regions of the ectoderm (Figure S4B-D). Next, we imposed a pattern of viscosity, with viscosity being highest near the margin and lowest at the animal pole (Figure 4E and S4E). Interestingly, we found that the animal pole ectoderm undergoes faster thinning compared to more marginal regions (Figures 4G and S4F-G) and that this thinning behaviour matched the ectoderm tissue spreading behaviour observed experimentally during LME migration (Figures 4D and 4F). Together, these data indicate that differences in ectoderm tissue viscosity along its AnVeg axis can explain its differential thinning behaviour.

We have previously shown that in the early zebrafish embryo, the cellular organisation of blastoderm cells at the onset of doming (4.3 hpf) differs between the centre and margin of the blastoderm, with the margin forming a more cohesive tissue than the centre^25^. We further showed that these differences in cell connectivity led to the blastoderm centre being less viscous than the margin^27^. We thus hypothesized that similar variations in tissue cohesiveness might be responsible for the observed differences in ectoderm thinning and viscosity along its AnVeg axis. To challenge this hypothesis, we quantified the animal and lateral ectoderm cell fraction. Remarkably, we observed that the lateral ectoderm had a higher cell fraction (97% on average) than the animal ectoderm (87% on average) during the period of LME animal pole-directed migration (Figure 5A-B and Video S6), suggesting that the ectoderm tissue is less compact at the animal pole compared to lateral ectoderm. To determine whether the smaller cell fraction at the animal pole is due to fewer cell-cell contacts in this region, we analysed the percentage of cell perimeter occupied by cell-cell contacts in the animal and lateral ectoderm. In the animal ectoderm, the percentage of cell perimeter in contact with other ectoderm cells was considerably smaller than in the lateral ectoderm (Figure S5A-B and Video S7), suggesting that the animal ectoderm is less cohesive than lateral ectoderm during the period of LME animal migration. We have previously shown that cell-cell contact size is determined by the ratio of cortical tensions at the cell-cell to the cell-outside medium interfaces^27–30^. Thus, we hypothesized that the higher cell fraction and cell-cell contact engagement in the lateral compared to animal ectoderm is due to a lower ratio of these interfacial tensions. To test this, we quantified the contact angle between cells, as readout of the ratio of cortical tensions at the cell-cell to the cell-medium interfaces^27–30^, within the lateral and animal ectoderm when LME animal migration was underway (28 min from the onset of animal migration) (Figure S5C, D). In agreement with our results on cell fraction and cell-cell contact engagement, we found that the contact angle was significantly higher, and thus the ratio of cortical tensions at the cell-cell to the cell-medium interfaces lower, in the lateral compared to the animal ectoderm (Figure S5E, on average 0.79 in the lateral ectoderm, 0.89 in the animal ectoderm). Collectively, these findings suggest that animal ectoderm shows a lower cohesiveness than lateral ectoderm, and that this difference in tissue cohesion is responsible for the observed difference in tissue viscosity.

**Figure 5.**
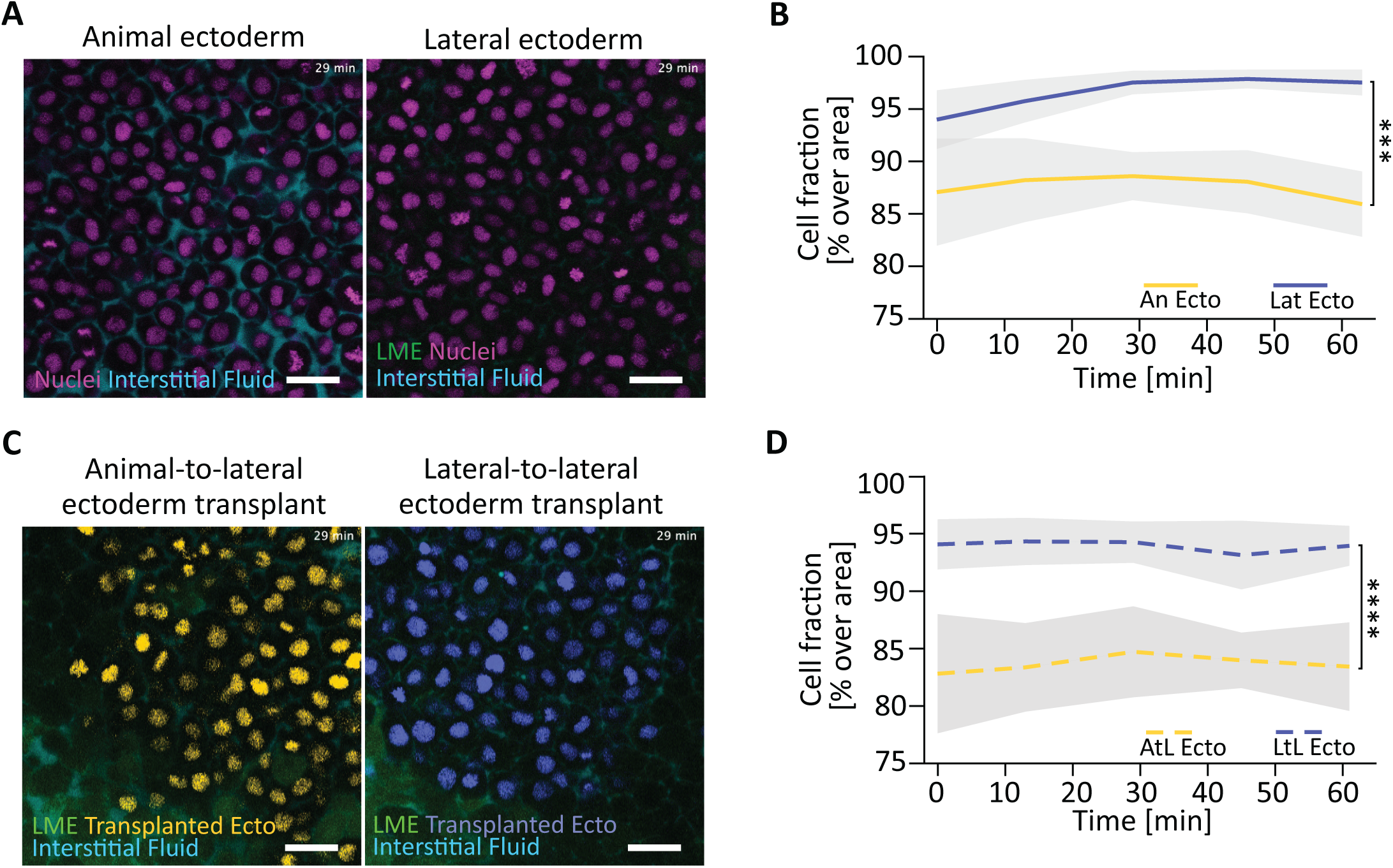
Lateral and animal ectoderm display different cell fractions. (A) Fluorescence images of animal (left) and lateral (right) ectoderm from wildtype (wt) *Tg(sebox::eGFP)* embryos injected with 35 pg *H2A-mCherry* mRNA (nuclei) at the 1-cell stage. At sphere/dome stage (4-4.3 hours post fertilization, hpf), embryos were injected with 1nl of 1 µg/µl 10.000 MW Alexa Fluor™ 647 to mark interstitial fluid. The images are maximum intensity projection of stacks of three 1 µm optical sections (Z step size, 1.5 µm). Green, eGFP (lateral mesendoderm (LME) cells); magenta, H2A-mCherry (nuclei); cyan, Alexa Fluor™ 647 dextran (interstitial fluid). Scale bar, 25 μm. See also Video S8. (B) Cell fraction in the animal and lateral ectoderm. Values are shown as mean (solid line) with standard deviation (SD, light grey area). Number of embryos: animal ectoderm (An Ecto, yellow), 5; lateral ectoderm (Lat Ecto, blue), 5. Statistical test, Two-way ANOVA: *** p<0.001. (C) Fluorescence images of animal-to-lateral (left) and lateral-to-lateral (right) ectoderm transplants. Animal (yellow) or lateral (blue) ectoderm cells from a wt donor *Tg(sebox::eGFP)* embryo injected with 35 pg *H2A-mCherry* mRNA to mark nuclei were transplanted at germ ring stage (5.7 hpf) into the lateral ectoderm of a stage-matched wildtype *Tg(sebox::eGFP)* host embryo injected with 1nl of 0.5 µg/µl 10.000 MW Alexa Fluor™ 647 dextran to mark interstitial fluid. The images are maximum intensity projection of stacks of three 1 µm optical sections (Z step size, 1.5 µm). Green, eGFP (LME cells); yellow, H2A-mCherry (nuclei, transplanted animal ectoderm); blue, H2A-mCherry (nuclei, transplanted lateral ectoderm); cyan, Alexa Fluor™ 647 dextran (interstitial fluid). Scale bar, 25 μm. See also Video S9. (D) Cell fraction in animal-to-lateral and lateral-to-lateral ectoderm transplants. Values are shown as mean (dashed line) with standard deviation (SD, light grey area). Number of embryos: animal-to-lateral ectoderm transplants (AtL Ecto, yellow), 11; lateral-to-lateral ectoderm transplants (LtL Ecto, blue), 12. Statistical test, Mixed-effect analysis: **** p<0.0001.

To establish a causative link between ectoderm tissue cohesion and LME animal migration, we first investigated whether, in our tissue transplantation experiments, the animal and lateral ectoderm tissues, when transplanted into the host embryos, kept their characteristic differences in tissue cohesion. Quantifying the cell fraction in lateral-to-lateral and animal-to- lateral transplants showed that the transplanted animal ectoderm (heterotypic transplantation) had a smaller cell fraction than the transplanted lateral ectoderm (homotypic transplantation) (Figure 5C-D and Video S8), similar to the situation in untransplanted embryos. To test whether the differences in tissue cohesion between animal and lateral ectoderm are decisive for their influence on LME animal migration, we increased cell contractility in the animal ectoderm by overexpressing constitutively active (CA) RhoA, previously shown to increase tissue cohesion and viscosity^25^, and we decreased cell contractility in lateral ectoderm by overexpressing CAMypt prior to performing our transplantation experiments (Figure 6A-B and Video S9). Remarkably, in the heterotypic transplantations (animal-to-lateral ectoderm transplantations), animal ectoderm overexpressing CARhoA transplanted into the lateral ectoderm lost its ability to block animal pole-directed LME migration. Conversely, in the homotypic transplantations (lateral-to-lateral ectoderm transplantations), LME cells encountering a less-contractile lateral ectoderm overexpressing CAMypt were hindered in their animal-pole-directed migration (Figures 6C-D and S5F-G). Together, these data show that the differences in ectoderm cohesion along the AnVeg axis are responsible for the different activities of animal and lateral ectoderm in modulating LME animal pole migration.

**Figure 6.**
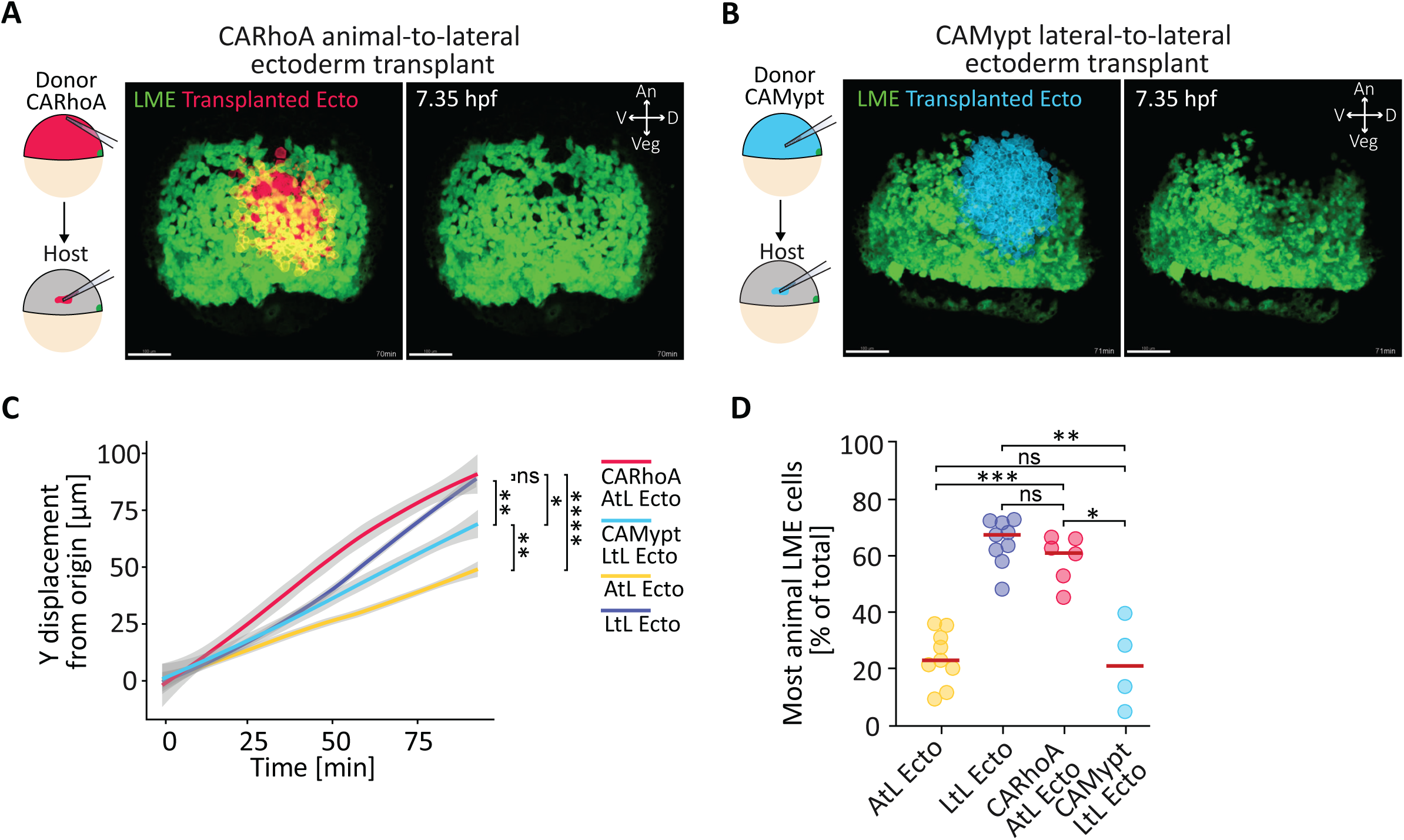
Modulation of cell contractility in the ectoderm reverses the effect of animal and lateral ectoderm on lateral mesendoderm migration (**A,B**) Constitutively active (CA) RhoA (CARhoA) overexpressing animal-to-lateral and CAMypt overexpressing lateral-to-lateral ectoderm transplant assay. Animal ectoderm cells (red, A) and lateral ectoderm cells (light blue, B) from a *Tg(sebox::eGFP)* donor embryo injected with either 1-2 pg *CARhoA* mRNA (A) or 75 pg *CAMypt* mRNA (B) together with 10-15 pg *H2B- mKO2k* and 100 pg *membrane RFP* mRNAs to mark nuclei and plasma membrane, respectively, were transplanted at germ ring stage (5.7 hours post fertilization, hpf) into the lateral ectoderm of a stage-matched wildtype *Tg(sebox::eGFP)* host embryo injected with 10- 15 pg *H2B-mKO2k* mRNA (nuclei). After transplantation, lateral mesenoderm (LME) cells encountering the transplanted ectoderm (for definition, see Figure S4 and Methods) were tracked for ∼92 min (∼2 min 14 sec frame rate, t0 ∼6.2 hpf). On the left, schematic representation of the experimental setup. On the right, fluorescence images of LME cells and transplanted CARhoA overexpressing animal (red, A) or CAMypt overexpressing lateral (light blue, B) ectoderm (lateral view) from representative timelapses ∼70 min after the start of LME migration (∼7.35 hpf). Green, eGFP (LME cells); red, membrane RFP (transplanted CARhoA overexpressing animal ectoderm); light blue, membrane RFP (transplanted CAMypt overexpressing lateral ectoderm). Scale bar, 100 μm. An, animal; Veg, vegetal; D, dorsal; V, ventral. See also Video S11 and S12. (C) Displacement of LME cells towards the animal pole over time in transplanted embryos. The displacement along the animal-vegetal (AnVeg) axis was quantified as cumulative Y-displacement from the origin of migration (Y-axis) and displayed as smooth plot. Solid line represents the mean, grey ribbon displays confidence interval. Number of cells: 292 from 6 CARhoA overexpressing animal-to-lateral ectoderm transplants (CARhoA AtL Ecto, red); 162 from 4 CAMypt overexpressing lateral-to-lateral ectoderm transplants (CAMypt LtL Ecto, light blue); 327 from 9 animal-to-lateral ectoderm transplants (AtL Ecto, yellow); 327 from 9 lateral-to-lateral ectoderm transplants (LtL Ecto, blue). Datasets for AtL Ecto and LtL Ecto correspond to data shown in Figure 3. Statistical test on the final Y-displacement, Mann-Whitney test: ns not significant, * p<0.05, ** p<0.01, **** p<0.0001. See also Figure S4 and S5. (D) Percentage of most animally-migrating LME cells in transplanted embryos. Tracked LME cells were scored as most animally-migrating at the end of the tracked time (∼92 min, 7.7 hpf) if they were within a 100 µm-wide bin (along the AnVeg axis) from the most animally-located LME cells that had not encountered the transplanted ectoderm (see also Figure S4). Red line represents the median. Number of embryos: CARhoA overexpressing animal-to-lateral ectoderm transplants (CARhoA AtL Ecto, red), 6; CAMypt for lateral-to-lateral ectoderm transplants (CAMypt LtL Ecto, light blue), 4; animal-to-lateral ectoderm transplants (AtL Ecto, yellow), 9; lateral-to-lateral ectoderm transplants (LtL Ecto, blue), 9. Datasets for AtL Ecto and LtL Ecto correspond to data shown in Figure 3. Statistical test, Mann-Whitney test: ns not significant, * p<0.05, ** p<0.01,*** p<0.001. See also Figure S5.

### BMP defines the non-permissiveness of the animal ectoderm to LME migration

To uncover the molecular pathway patterning tissue cohesion and viscosity within the ectoderm, we analysed how gene expression differs between lateral and animal ectoderm during LME migration. To this end, we obtained pure populations of either animal or lateral ectoderm cells by expressing a photoactivatable (PA) version of the fluorescent protein mCherry1 in wildtype (wt) *tg(Sebox::eGFP)* embryos and then selectively activating it in either the animal or the lateral ectoderm between 6 hpf and 7.5 hpf using a confocal microscope. After photoactivation, we isolated mCherry1-positive cells using fluorescence-activated cell sorting (FACS) and analysed them by bulk RNA sequencing (Figure S6A-B). We found that 54 and 81 genes were predominantly expressed in the animal and lateral ectoderm, respectively (adjusted p-value < 0.05) (Figure S6B and Table S1). An initial analysis of those genes showed that the lateral ectoderm predominantly expressed non-nuclear components of several key signalling pathways, including extracellular effectors of the Wnt and BMP pathways (*wnt8a*, *wnt5b*, *wnt11f2* and *chrd* Figure S6B and Table S1). Given that BMP signalling has previously been shown to be required for zebrafish LME cell convergence movements^17, 19^, we asked whether the transition of LME cells from animal pole-directed migration to the tumbling phase might also depend on BMP signalling. To this end, we determined how depletion of the BMP ligand Bmp2b affects LME migration. Interestingly, we observed that in *bmp2b* morphant embryos, both LME and ventral mesendoderm failed to transit into their tumbling phase and, instead, continued migrating towards the animal pole until they collided with the prechordal plate (Figure 7A, Video S11). To assess how BMP functions in this process, we performed ectoderm tissue transplantation experiments, replacing the lateral ectoderm of wt embryos with the animal ectoderm of *bmp2b* morphant embryos (Figure 7B, Video S12). This showed that the transplanted BMP-depleted animal ectoderm failed in slowing down LME animal pole-directed migration (Figure 7C-D, Figure S6C-D), indicating that BMP signalling is required for the activity of animal ectoderm in slowing down animal pole-directed LME migration. Finally, we asked whether BMP signalling functions in this process by lowering animal ectoderm tissue cohesion. To this end, we quantified the cell fraction in BMP-depleted animal-to-lateral transplants and found that the cell fraction in transplanted BMP-depleted animal ectoderm (heterotypic transplantation) was increased to a level typically found for transplanted lateral ectoderm in homotypic transplantations (Figure 7E). Together, these data indicate that BMP signalling reduces cell cohesion and, thus, tissue viscosity in the animal ectoderm, thereby rendering this tissue non-permissive for LME animal pole-directed migration.

**Figure 7.**
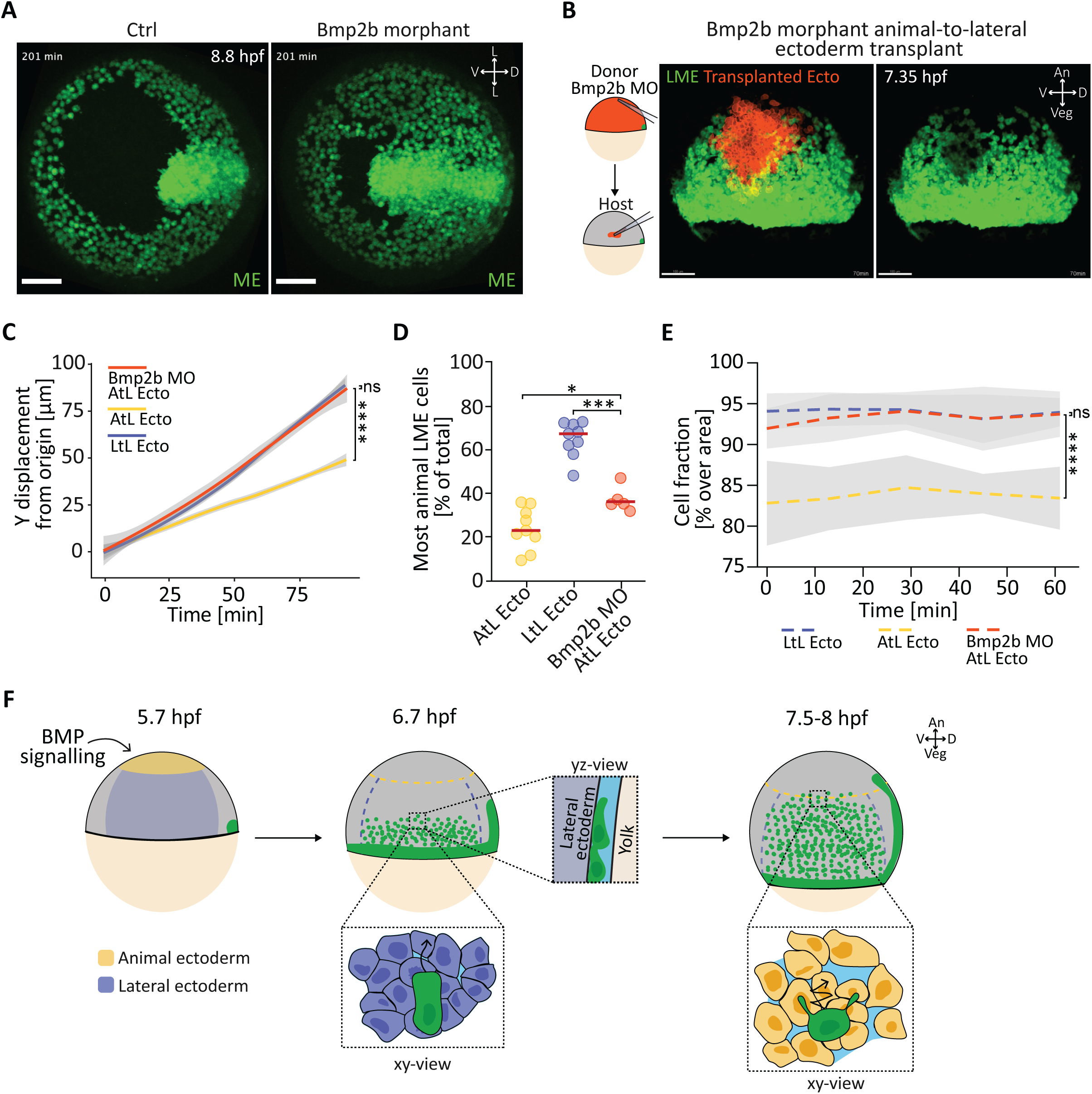
BMP signalling is needed for animal ectoderm blocking animal pole-directed lateral mesendoderm migration. (A) Fluorescence images of mesendoderm (ME) cells (animal view) in *Tg(sebox::eGFP)* embryos injected with 1.5 ng control (Ctrl) or 1.5 ng *bmp2b* (Bmp2b morphant) *morpholinos* (MO), shortly before the onset of ME convergence movements in the control embryo. Green, eGFP (ME cells). Scale bar, 100 μm. L, lateral; D, dorsal; V, ventral. See also Video S13. (B) *Bmp2b* morphant animal-to-lateral ectoderm transplant assay. Animal ectoderm cells (orange) from a *Tg(sebox::eGFP)* donor embryo injected with 1.5 ng *bmp2b* MO together with 15 pg *H2B-mKO2k* and 100 pg *membrane RFP* mRNAs to mark nuclei and plasma membrane, respectively, were transplanted at germ ring stage (5.7 hours post fertilization, hpf) into the lateral ectoderm of a stage-matched wildtype *Tg(sebox::eGFP)* host embryo injected with 10- 15 pg *H2B-mKO2k* mRNA (nuclei). After transplantation, lateral mesendoderm (LME) cells encountering the transplanted ectoderm (for definition, see Figure S4 and Methods) were tracked for ∼92 min (∼2 min 14 sec frame rate, t0 ∼6.2 hpf). On the left, schematic representation of the experimental setup. On the right, fluorescence images of LME cells and transplanted *bmp2b* MO animal ectoderm (orange) (lateral view) from a representative timelapse movie ∼70 min after the start of LME migration (∼7.35 hpf). Green, eGFP (LME cells); orange, membrane RFP (transplanted *bmp2b* MO animal ectoderm). Scale bar, 100 μm. An, animal; Veg, vegetal; D, dorsal; V, ventral. See also Video S14. (C) Displacement of LME cells towards the animal pole over time in transplanted embryos. The displacement along the animal-vegetal (AnVeg) axis was quantified as cumulative Y- displacement from the origin of migration (Y-axis) and displayed as smooth plot. Solid line represents the mean, grey ribbon displays confidence interval. Number of cells, 199 from 5 *bmp2b* MO animal-to-lateral ectoderm transplants (Bmp2b MO AtL Ecto, orange); 327 from 9 animal-to-lateral ectoderm transplants (AtL Ecto, yellow); 327 from 9 lateral-to-lateral ectoderm transplants (LtL Ecto, blue). Dataset for AtL Ecto and LtL Ecto corresponds to data shown in Figure 3. Statistical test on the final Y-displacement, Mann-Whitney test: ns not significant, **** p<0.0001. See also Figure S4 and S6. (D) Percentage of most animally-migrating LME cells in transplanted embryos. Tracked LME cells were scored as most animally-migrating at the end of the tracked time (∼92 min, 7.7 hpf) if they were within a 100 µm-wide bin (along the AnVeg axis) from the most animally-located LME cells that had not encountered the transplanted ectoderm (see also Figure S4). Scatter dot plot shows the mean of each embryo. Red line represents the median. Number of embryos: 5 for *bmp2* MO animal-to-lateral ectoderm transplants (Bmp2b MO AtL Ecto, orange); 9 for animal-to-lateral ectoderm transplants (AtL Ecto, yellow); 9 for lateral-to-lateral ectoderm transplants (LtL Ecto, blue). Datasets for AtL Ecto and LtL Ecto correspond to data shown in Figure 3. Statistical test, Mann-Whitney test: ns not significant, * p<0.05, *** p<0.001. See also Figure S4 and S6. (E) Cell fraction in *bmp2b* morphant (injected with 1.5 ng *bmp2b* MO) animal-to-lateral ectoderm transplants. Values are shown as mean (dashed line) with standard deviation (SD, light grey area). Number of embryos: 6 for *bmp2* MO animal-to-lateral ectoderm transplants (Bmp2b MO AtL Ecto, orange); 11 for animal-to-lateral ectoderm transplants (AtL Ecto, yellow); 12 lateral-to-lateral ectoderm transplants (LtL Ecto, blue). Datasets for AtL Ecto and LtL Ecto correspond to data shown in Figure 5. Statistical test, Mixed-effect analysis: ns not significant, **** p<0.0001. (F) Schematic of ectoderm-dependent modulation on animal pole-directed migration of LME cells: from germ ring stage (5.7 hpf) onwards animal ectoderm becomes non-permissive for animal pole-directed LME migration. This function of animal ectoderm in blocking animal pole-directed LME migration depends on BMP2b expression within the animal pole ectoderm. An, animal; Veg, vegetal; D, dorsal; V, ventral.

## Discussion

Our work shows that in the early zebrafish gastrula, the ectoderm plays a decisive role in LME migration by functioning as a substrate controlling LME cell migration. In gastrulation, mesendoderm cells, upon internalisation, typically become more mesenchymal and migratory to move away from their site of internalisation and migrate to their final destination^28^. We now show that in zebrafish, internalised mesendoderm cells in lateral regions of the gastrula use the overlying ectoderm as their prime substrate for migration. This choice of a cellular substrate instead of ECM is likely due to zebrafish embryos showing distinct ECM accumulations only towards the end of gastrulation^7, 8^, reminiscent of the situation in *Drosophila*, where ECM is not yet present during mesoderm migration and mesoderm cells move dorsolaterally using the ectoderm as their substrate^29^. In contrast, ECM accumulation is already clearly recognizable in frog, chick and mouse embryos at the onset of gastrulation^28, 30–34^. Consequently, in *Xenopus*, upon involution, the leading edge mesendoderm and prechordal mesoderm undergo collective migration towards the animal pole using fibronectin fibrils formed at the ectodermal blastocoel roof as their substrate for migration^28, 32^. Likewise, in chick and mouse, mesoderm cells undergo collective mesenchymal migration between the ectoderm and extraembryonic endoderm using basal lamina as their substrate for migration^28, 30, 31, 33, 34^. This suggests that the role of ECM for mesoderm migration can vastly differ among different model organisms.

How LME cells migrate independently of ECM in zebrafish is not yet entirely clear. Previous studies *in vitro* and *in vivo* have suggested that cells undergo ECM-independent migration using cell-cell adhesion or unspecific friction with surrounding structures^4, 35, 36^. Cell-cell adhesion as a mechanism for cells to generate traction force for migration has been, for instance, proposed in *Drosophila* to be used by both border cells during oogenesis^37, 38^ and mesoderm cells during gastrulation^29^. It is thus conceivable that LME cells use cell-cell adhesion for migrating on the ectoderm, although the precise role of cell-cell adhesion for traction force generation by LME cells remains to be explored. Alternatively, LME cells might undergo migration by generating unspecific friction with their surrounding environment. Such migration mode has been proposed to be particularly relevant for cells placed in spatial confinement^4, 35, 36^. Consistent with the possibility of LME cells migrating by unspecific adhesion, LME cells during their migration are spatially confined between the yolk cell and ectoderm. However, our observation that reducing this spatial confinement by increasing the space between yolk cell and ectoderm in Mannitol-injected embryos had no major effect on LME migration, argues against LME cells undergoing unspecific friction-mediated migration.

Our findings also show that while the majority of LME cells preferentially use the ectoderm as their substrate for migration, the lateral endoderm cells appear to be equally subdivided between those preferring either the ectoderm or the yolk cell. Interestingly, we also found that lateral endoderm cells migrating on the ectoderm show more directed and persistent movements than those using the yolk cell as their substrate for migration. It has previously been shown that Nodal signalling promotes the random motion of endoderm cells through Prex1-dependent activation of Rac1^39^. Given that the YSL is an important source of Nodal ligands before gastrulation^40–42^, it is an intriguing possibility that Nodal signals emanating from the YSL trigger random motion of endoderm cells moving on the yolk cell. Yet, it still needs to be investigated how lateral mesoderm and endoderm cells choose their substrate for migration, and how such choice of substrate influences their migration directionality and persistence.

One major function of the ectoderm in zebrafish appears to determine how far LME cells can migrate towards the animal pole before switching into their tumbling phase. In the absence of confinement, there are three main means by which the substrate can affect cell migration^1^: migrating cells can respond to differences in substrate topology, as demonstrated, e.g. in the zebrafish developing optic cup^43^, to gradients of ECM- or cell-bound components, as found e.g. for immune cells haptotaxis^44, 45^, or to differences in substrate stiffness, as demonstrated e.g. for neural crest cells migration in *Xenopus* ^46^. Our observations suggest that differences in ectoderm viscosity/stiffness are responsible for LME cells switching into their tumbling phase when getting closer to the animal pole, where ectoderm viscosity is lower. This suggests that LME cells respond to differences in substrate stiffness. However, they do not seem to respond to a stiffness gradient, as they can migrate animally on ectoderm tissue with uniform high tissue viscosity upon overexpression of CARhoA. Rather they seem to respond to a stiffness threshold, which determines whether LME cells can undergo directed migration or tumble. This is similar to neural crest cells in *Xenopus*, which start migrating when the head mesoderm reaches a specific stiffness^47^.

How tissue viscosity/stiffness affects the migration mode of LME cells is still unclear, but mechanosensation of lateral mesodermal and/or endodermal cells is likely to be involved. Previous studies have shown that Cxcl12b/Cxcr4a signalling couples lateral mesoderm and endoderm migration and that, in the absence of such signalling, lateral endoderm cells do not switch into a tumbling phase and, instead, continue migrating towards the animal pole^13, 14, 48^. Therefore, a likely scenario is that mesoderm, but not endoderm, can distinguish lateral from animal ectoderm through a mechanosensitive mechanism. The nature of this mechanism is still unknown, but given that LME cells are likely to use cadherin-mediated cell-cell adhesion to adhere to and migrate along the ectoderm, mechanosensation of the E-cadherin adhesion complex^49, 50^ might be involved.

While the central role of tissue material properties in various key developmental processes, such as migration of chick hindgut endoderm^51^, *Drosophila* germband extension^52^, *Tribolium* serosa expansion^53^, zebrafish doming^25^ and body axis elongation^54^, becomes increasingly clear, questions remain as to the molecular and cellular mechanisms regulating these properties. We have previously shown that between sphere and dome stage (4-4.3 hpf), the blastoderm undergoes a transient fluidization at its centre but not the margin^25^. We further showed that this lack of tissue fluidization at the margin was due to Wnt/planar cell polarity (Wnt/PCP) signalling at the blastoderm margin increasing cell cohesion^25^. Our study points to yet another signalling pathway, BMP signalling, being involved in modulating tissue viscosity. BMP signalling has previously been shown to control the directionality of LME convergence movements by lowering cadherin-mediated cell-cell adhesion in a concentration-dependent manner^19^. This, together with our finding that cell cohesion is lower in animal compared to lateral ectoderm, suggests that one possible pathway through which BMP reduces tissue viscosity in the animal ectoderm is by lowering cadherin-mediated cell- cell adhesion. Interestingly, whereas BMP forms a gradient along the dorsoventral (DV) axis of the gastrula, with peak levels at the ventral side, ectoderm tissue viscosity differs along the AnVeg axis of the ectoderm. Why BMP lowers ectoderm tissue cohesion only in the animal but not lateral ectoderm is not yet clear, but it is conceivable that other signalling pathways, specifically active within the lateral ectoderm, such as the Wnt/PCP and canonical Wnt pathways, might counteract the activity of BMP signalling in lowering tissue viscosity there. In line with this possibility are recent reports showing that BMP target gene expression can be modulated by the co-activation of other signalling pathways, such as the Nodal and Fgf pathways^55^.

The interaction between the different germ layers represents a key factor determining embryo morphogenesis during gastrulation^56^. While there is increasing evidence for the mesendoderm to affect ectoderm morphogenesis^57–59^, much less is known about how ectoderm feeds back on mesendoderm morphogenesis. By showing that ectoderm tissue material properties control the migratory properties of LME cells, we identify a critical role of ectoderm not only in providing a substrate for mesendoderm cell migration but also instructing the migration mode by which LME cells drive mesendoderm morphogenesis.

## STAR Methods

### KEY RESOURCES TABLE

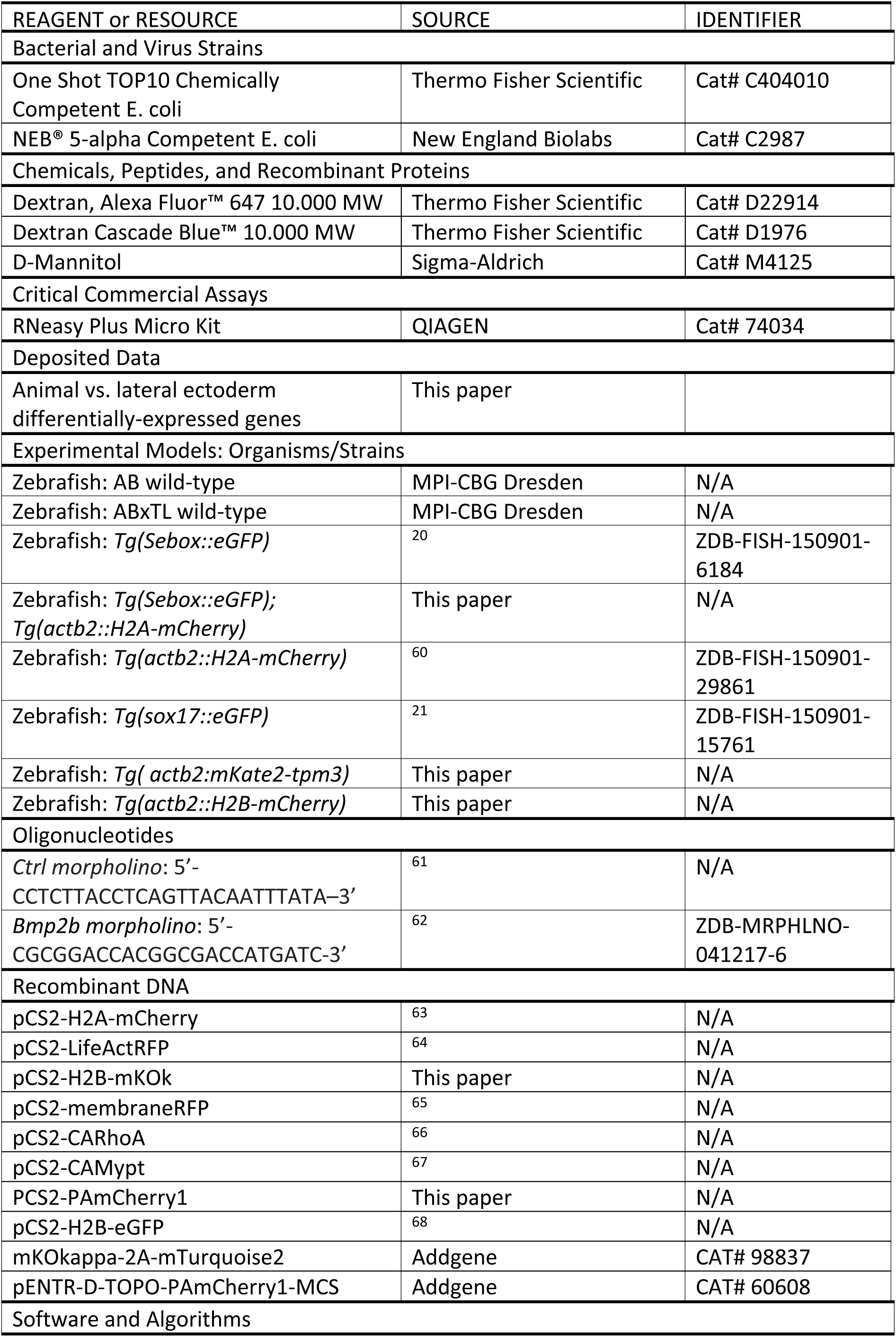
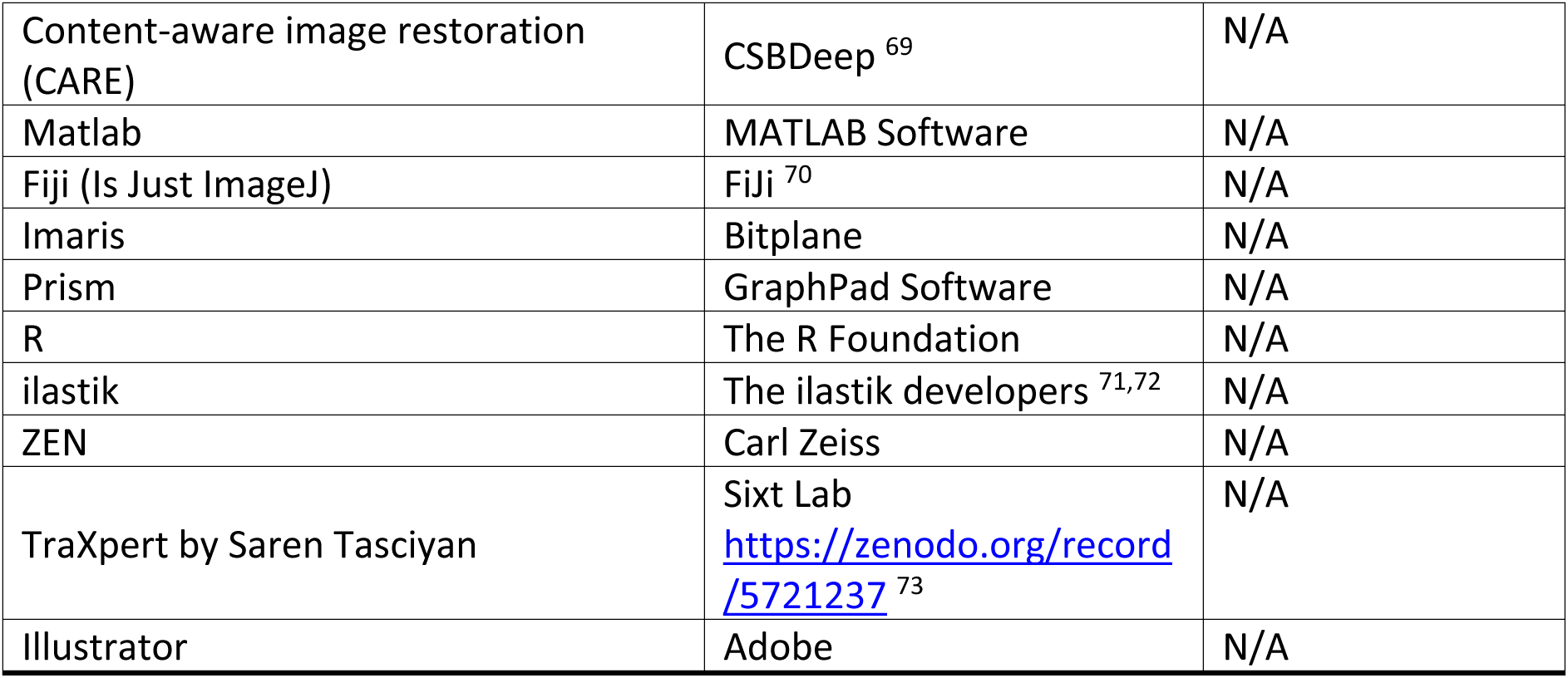

### CONTACT FOR REAGENT AND RESOURCE SHARING

Further information and requests for resources and reagents should be directed to Carl- Philipp Heisenberg.

### EXPERIMENTAL MODEL AND SUBJECT DETAIL

#### Fish

Zebrafish (*Danio rerio*) were housed in 28C water (pH 7.5 and conductivity 400mS) with a 14 h on/10 h off light cycle. All zebrafish husbandry and breadings were performed in the zebrafish facility at IST Austria under standard conditions according to local regulations, and all procedures were approved by the Ethics Committee of IST Austria regulating animal care and usage. Embryos were raised in E3 medium and kept at 25-31C until start of the experimental procedures (4 hpf - 5.7 hpf). After dechorionation and for all experimental procedures preceding live imaging, embryos were kept in Danieau buffer. Staging of the embryos was done according to Kimmel et al^74^.

### METHOD DETAILS

#### Plasmids

All DNA plasmids were extracted and purified using Monarch® Plasmid Miniprep Kit (QIAGEN) or EndoFree Plasmid Maxi kit (QIAGEN) following the manufacturer instructions. The Gateway technology^75, 76^ was used to generate *pCS2-H2B-mKOk* as follows: the coding sequence for H2B and mKOk were amplified from *pCS2-H2B-eGFP* and from *mKOkappa-2A-mTurquoise2*, respectively. The PCR products were then recombined with *pDONR221* (Lawson#208) for H2B and with *pDONR P2r-P3* (Lawson#211) for mKOk, and subsequently with *pCSDest2* (Lawson#404). The *pCS2-PAmCherry1* plasmid was generated starting from *pCSDest2* (Lawson#404). The coding sequence for PAmCherry1 was PCR-amplified from *pENTR-D-TOPO-PAmCherry1-MCS*. The fragments were subsequently ligated using NEBuilder® HiFi DNA Assembly Cloning Kit (New England Biolabs) following the manufacturer instructions.

#### Transgenic zebrafish line generation

*Tg(actb2:mKate2-tpm3)* and *Tg(actb2::H2B-mCherry)* transgenic lines ubiquitously expressing mKate2-tagged tpm3 and mCherry-tagged H2B, respectively, were generated using the Tol2/Gateway technology^75, 76^.

For *Tg(actb2:mKate2-tpm3)*, the coding sequence of *tpm3* (NCBI reference sequence: NM_201492.2) was amplified using gene specific primers with additional Gateway recombination arms (5ʹ- GGGGACAGCTTTCTTGTACAAAGTGGCTATGGCCGGATCAAACAGC -3ʹ and 5ʹ- GGGGACAACTTTGTATAATAAAGTTGCTTAGAAACTATTGAGCTCGCTCAG -3ʹ) from cDNA library of sphere stage wild-type Tübingen embryos. The PCR product was recombined with *pDONR P2r-P3* (Lawson#211) and the resulting entry clone was recombined with *pDestTol2pA2* (Chien#394), *p5E β-actin* promoter (Chien#229) and *pME mKate2* to create *pTol2-β-actin:mKate2-tpm3*. The *pTol2* vector was co-injected with mRNA encoding the transposase (Invitrogen) into 1 cell-stage wild-type TL embryos. Individual positive carriers were selected and out-crossed with wild-type TL fish for stable single-copy genetic integration.

For *Tg(actb2::H2B-mCherry)*, the coding sequence of H2B was amplified from *pCS2-H2B-eGFP*. The PCR product was recombined with *pDONR P2r-P3* (Lawson#211) and the resulting entry clone was recombined with *pDestTol2pA2* (Chien#394), *p5E β-actin* promoter (Chien#229) and *p3E mCherry+pA* (Chien#388) to create *pTol2:actb2-H2B-mCherry_polA*. The *pTol2* vector was co-injected with mRNA encoding the transposase (Invitrogen) into 1 cell-stage wild-type AB embryos. Individual positive carriers were selected and out-crossed with wild-type AB fish for stable single-copy genetic integration.

*Tg(Sebox::eGFP); Tg(actb2::H2A-mCherry)* transgenic line was generate by crossing the pre-existing *Tg(Sebox::eGFP)* and *Tg(actb2::H2A-mCherry)* transgenic lines.

#### mRNA, Morpholino, Mannitol and dextran injections

mRNA transcription was performed using the SP6 mMessage mMachine Kit (Thermo Scientific). Glass capillaries (30-0020, Harvard Apparatus) were pulled using a needle puller (P-97, Sutter Instruments) and mounted on a microinjection system (PV820, World Precision Instruments). Injections at 1-cell and 64-128-cell stages were performed as described^77^. The following mRNAs were injected: 35-50 pg *H2A-mCherry*, 10-15 pg *H2B-mKOk*, 100 pg *membrane-RFP*, 1-2 pg *CARhoA*, 75 pg *CAMypt* and 125 pg *PAmCherry*. For labelling F-actin, 7.5 pg *LifeactRF*P were injected into a single blastomere at 64-128-cell stage. For gene knock- down experiments, the following morpholinos were used in this study: 1.5ng *bmp2b* MO (5’- CGCGGACCACGGCGACCATGATC-3’), 1.5 ng Ctrl MO (*human β-globin* MO 5’-CCTCTTACCTCAGTTACAATTTATA–3’). For Mannitol injection, 1 nl of 350 mM Mannitol with 1.25 µg/µl 10.000 MW Alexa Fluor™ 647 dextran was injected in the interstitial space of sphere/dome stage (4-4.3 hpf) embryos. For labelling the interstitial fluid of embryos, 1nl of either 1 µg/µl 10.000 MW Alexa Fluor™ 647 dextran or 1.5 µg/µl 10.000 MW Cascade Blue™ dextran was injected in the interstitial space of sphere/dome stage (4-4.3 hpf) embryos. To mark the interstitial fluid of embryos after tissue transplantation, 1nl of 0.5 µg/µl 10.000 MW Alexa Fluor™ 647 dextran was injected in the interstitial space of germ ring (5.7 hpf) embryos.

#### Yolk peeling assay

The yolk membrane was manually peeled from the blastoderm of *Tg(sebox::eGFP)* embryos at 65-70% epiboly (7-7.5 hpf) using forceps, and both embryo and yolk membrane were immediately fixed in 4% PFA. Both blastoderm and yolk membrane were then stained with DAPI (nuclei) before imaging.

#### Deep-cell transplantations

At germ ring stage (5.7 hpf), both host and donor embryos were transferred to Danieau buffer and prepared for transplantation. Using a spike fire-polished transplantation needle with a 20-μm inner diameter (Biomedical Instruments) attached to a syringe system via silicone tubing, 100-200 ectoderm cells were taken from the donor embryos and placed into the lateral ectoderm of the host embryo. Transplanted host embryos were then mounted for upright and inverted imaging, depending on the experimental assay.

#### Sample preparation for live imaging

Embryos were dechorionated and mounted either in in 0.65% LMP agarose and put into prepared agarose molds (2%) for up-right imaging or in 0.5% low melting point (LMP) agarose (Invitrogen) on glassbottom dishes (MatTek) for inverted imaging. For all live imaging, the temperature during image acquisition was set to 28.5C.

#### Imaging setups for live and fixed imaging

Up-right imaging was performed using either a Zeiss 800 confocal microscope equipped with a Plan-Apochromat 20X/1.0 W objective or a Leica SP8 confocal microscope equipped with a HC FUOTAR L 25x/0.95 W objective. For live imaging, Z-stacks of 58-255 µm with a Z step-size of 1.5 µm and a Z-thickness of 2 µm were acquired every 2-9 min. Inverted imaging was performed using either a Zeiss 800 confocal microscope equipped with a Plan-Apochromat 40X/1.2 W objective or a Leica SP8 confocal microscope equipped with a HC PL APO 40x/1.10 W objective. For live imaging, Z-stacks of 40-75 µm with a Z step-size of 1.5 µm and a Z-thickness of 1 µm were acquired every 2-6 min.

#### LME displacement and delta displacement

Time lapse movies of the embryo lateral side were acquired with a time resolution of approximately 6.2 min. Subsequently, a cross-section view of the embryo was created from a 20 µm-wide volume (Video S1, dashed lines) from the middle of the lateral view using “Reslice /” and “Average Z Projection” plugins in Fiji (Is Just ImageJ) (FiJi). The cross-section view was then used to track the distance covered along the embryo curvature by the front edge of the LME every 23 min, starting from the origin of migration (6.2 hpf) for at least 115 min. The delta displacement was computed as difference of each time point from the previous one. Data were plotted using Prisms (GraphPad).

#### Protrusion directionality

To compute protrusion directionality, the centre and radius of the embryo were determined by approximating the surface shape of the embryo as a sphere and then fitting the embryo surface coordinates obtained by using Spots feature in Imaris (Bitplane). For each cell, the coordinates of the tip of each visible actin protrusions and the cell centre were then obtained using Spots feature in Imaris (Bitplane). Using these coordinates, the radial distance was computed as the difference between the protrusion tip position relative to the embryo centre and the cell centre positions. Value above 0 were defined as ectoderm-directed protrusion, while value below 0 as yolk-directed protrusions. Protrusion lengths was quantified as distance of the protrusion tip from the cell centre. Data were plotted as percentage over total number of protrusions and analysed using Prisms (GraphPad).

#### Single cell tracking

For tracking of single cells, timelapse movies were acquired with a temporal resolution of approximately 2 min. If necessary, the spatial resolution of the acquired timelapse images was restored to 1024x1024 pixel after acquisition using Content-aware image restoration (CARE). The nuclear signal (H2B-mCherry or H2B-mKok) was segmented using Surfaces and tracks were computed using the Autoregressive Motion algorithm in Imaris 9 (Bitplane). Each track was subsequentially manually checked and corrected if necessary. In case of cell division, the track of one daughter cell was deleted. If not differently stated, cell migration data were then imported in TraXpert for subsequent statistical analyses and plots creation. Smooth plots were generated with ggplot’s geom_smooth function, which uses local polynomial regression fitting.

For determining the directness of LME and lateral endoderm cell migration, the tracks were exported and the overall direction of the migration of all cells was calculated using a custom script. The mean alignment of the cell movement with respect to this reference axis was computed as the cosine of the angle between the direction of cell motion at a particular time point and the established reference axis. For analysing the time LME cells spent on the ectoderm, the nuclei of each cell ware used as reference. At each time point, LME or lateral endoderm cells were scored as being placed on the ectoderm when the nucleus was in close proximity with the ectoderm layer. Close proximity was define being placed at a distance as equal or less than half nuclear diameter with no o little interstitial fluid (10.000 MW Alexa Fluor™ 647 dextran) in between ectoderm layer and LME or lateral endoderm nuclei. Data were plotted as percentage of total migration time or as relative frequency using Prisms (GraphPad). For determining the correlation between animal displacement and the time LME cells spent on the ectoderm, the total displacement along the Y axis (Y displacement, corresponding to the AnVeg axis) from the origin and the total time spent of the ectoderm were extracted from the tracking data of each cell. Statistical analysis and plotting of data were done using Prisms (GraphPad). “First rows after ingression” and “First ingressing cells” refer to LME cells or lateral endoderm cells located in the first 2-3 cell rows at the start of the tracking (6.2 hpf). In the case of transplantation experiments, LME cells were selected only if they were located in an area vegetal to the transplanted ectoderm, in an area that was 1-2 cell diameters smaller in width on the X axis (DV axis) on both sides in comparison to the transplant itself (see Figure S4). “All” refers to LME cells or lateral endoderm cells located in a 300 x 200 µm rectangular area with the front end of this area given by the most animally- located cells at the end of the tracking (7.7 hpf). “Most animal” refers to LME cells or lateral endoderm cells located in a 300 x 100 µm rectangular area with the front end of this area given by the most animally-located cells at the end of the tracking (7.7 hpf). In transplanted embryos, tracked LME cells were score as “Most-animally migrating cells” if they were within a 100 μm-wide bin on the Y axis (AnVeg axis) set following the final position of the most- animal LME cells that have not encountered the transplanted ectoderm (See Figure S4).

#### Ectoderm thickness

Time lapse movies were acquired of either the embryo animal pole or the lateral side with a time resolution of approximately 6.2 min. Subsequently, a cross-section view of the embryos was created from a 20 µm-wide volume from the middle of either the lateral view or the animal pole view, the latter spanning from the lateral to lateral, using the “Reslice /” and “Average Z Projection” plugins in Fiji (Is Just ImageJ) (FiJi). The cross-section views were then used to create a binary mask of the embryo, the LME and the interstitial fluid using ilastik. To quantify the ectoderm thickness from the binary masks, kymographs were created oriented along the inside-out axis of the tissue at 100 μm intervals, starting either from the animal pole or 100-150 µm animal of the embryo equator. For animal and lateral ectoderm thickness, bins were defined as either lateral ectoderm, in case the ectoderm was in contact with the LME at least for one timepoint during the imaging period, or animal ectoderm in case the ectoderm was not in contact with LME during the entire imaging period. Statistical analyses and plots were done using Prisms (GraphPad).

#### Cell fraction

Time lapse movies were acquired from either the embryo animal pole or the lateral side. In the case of untransplanted embryos, for each time point 2 not overlapping z-planes were chosen in the middle of the cell layer closest to the yolk membrane. In case of the animal to lateral (heterotypic) and lateral to lateral (homotypic) transplants, 1 to 4 not overlapping z-planes were chosen in the middle of the transplant. For each z-plane, the fluorescent signal of 10.000 MW Alexa Fluor™ 647 was used to create binary masks of the interstitial fluid using ilastik. The binary masks were then used to quantify the percentage of cell fraction over area using using Fiji (Is Just ImageJ) (FiJi). Statistical analyses and plots were done using Prisms (GraphPad).

#### Cell-cell contacts

Time lapse movies were acquired from either the embryo animal pole or the lateral side. The z-stacks containing the layer of ectoderm cells closest to the yolk were then used for the analysis. From each z-plane of the z-stacks, the fluorescent signal of 10.000 MW Cascade Blue™ dextran was used to create binary masks of the interstitial fluid using ilastik. The binary masks were then used to quantify the percentage of cell perimeter not in contact with interstitial fluid using Fiji (Is Just ImageJ) (FiJi). The cell perimeter was defined as the cell membrane (labelled with membrane RFP) in the z-plane having the widest XY cross-section area of the cell nucleus (labelled with H2A-mCherry). Statistical analyses and plots were done using Prisms (GraphPad).

#### Relative cell tension

Time lapse movies were acquired from either the embryo animal pole or the lateral side. The z-stacks containing the layer of ectoderm cells closest to the yolk were then used for the analysis. From each z-plane of the z-stacks, the fluorescent signal of 10.000 MW Cascade Blue™ dextran was used to create binary masks of the interstitial fluid using ilastik. The binary masks were then used to determine the presence or absence of interstitial fluid. Contact angles 8 were quantified between two ectoderm cells at the interphase with interstitial fluid using the angle tool in Fiji (Is Just ImageJ) (FiJi), with at least one of the two cells being in the z-plane with the widest XY cross-section area of the cell nucleus (labelled with H2A-mCherry). The contact angles were then converted in radiant. The relative cell tension α was then computed as

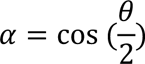

Statistical analyses and plots were done using Prisms (GraphPad).

#### Photoactivation of PAmCherry1 and FACS

To specifically mark animal or lateral ectoderm, *Tg(Sebox::eGFP)* embryos injected with *PAmCherry1* mRNA were mounted for up-right imaging and a 444 x 220 x 58.5 µm volume was photoactivated using a 405 nm laser either at the animal pole or on the lateral side. The photoactivation process was composed of 10 cycles of 28 sec per embryo in a time window of approximately 90 min from 6 to 7.5 hpf. After photoactivation, embryos were transferred in Ca^2+^-free Ringer solution and dissociated by gently pipetting using a P-1000 pipette to obtain a single-cell suspension. Per replicate, 5000 PAmCherry1+/eGFP - cells were sorted in 150 µl of RTL plus lysis buffer (RNeasy Plus Micro Kit, QIAGEN) with 1% of 2-mercaptoethanol, briefly vortexed, centrifuged in a small tabletop centrifuge and stored at –80 °C until RNA extraction.

#### RNAseq

Cells were subjected to RNA extraction using RNeasy Plus Micro Kit (QIAGEN) according to the manufacturer instructions. Complete cDNA synthesis and library preparation were performed using SMART-Seq v3 protocol^78^ with Nextera UDI adapters. Libraries were then quantified by qPCR (KAPA Biosysytems) and subjected to NovaSeq S4 XP 150-bp pair end sequencing on the NovaSeq 6000 platform, providing on average reads per sample of 92963813.5.

#### Transcriptome data processing and analysis

Reads of the same sample on different sequencing lanes were subject to quality checks with fastqc (0.11.7), before and after adapter and quality trimming with trimmomatic (version: 0.38; parameters: 2:30:10 LEADING:3 TRAILING:3 SLIDINGWINDOW:4:15 MINLEN:19). The processed reads were aligned and quantified using Salmon (1.1.0)^79^ in mapping based mode, with the --seqBias and --gcBias flags set in order to decrease the hexamer priming and GC content biases. For this quantification, an index created from the zebrafish reference genome (GRCz11) was used. The differential expression analysis was performed using the following libraries: DESeq2 (v. 1.30.1), apeglm (v. 1.12.0), tximport (v. 1.18.0), genefilter (v. 1.72.1) and GenomicRanges (v. 1.42.0). The expression of each gene was expressed as a transcript per million (TPM) value. Differential expression was determined without distinguishing between splice variants of the gene under study and using all protein-coding genes as transcriptome reference. Differentially expressed genes were defined using a cut-off of adjusted P-value <0.05. The statistical analysis and the calculation of the numbers of differentially expressed genes in the various sectors of the Volcano plot were calculated in R using the Enhancedvolcano (1.8.0) and biomaRt (2.46.3) libraries.

### SUPPLEMENTARY THEORY NOTE

As a minimal physical model of the differential thinning of the ectoderm layer, we use a fluid model with patterned viscosity. In previous work, a simple model for tissue fluidization at the onset of doming predicted the relative thickness change of the blastoderm as two tissues with different viscosities^25^. Here, we generalize this approach to the case of a continuously varying patterned viscosity and apply it to ectoderm morphodynamics in the early gastrula, i.e. a different tissue at a different time of development.

We model the ectoderm as an incompressible passive fluid that is extended by an active force *fA* at the lateral sides, enforcing a constant expansion speed *v*. We define a one-dimensional coordinate system in which the initial tissue is spread from *x* = −*L* to *x* = *L*, where *x* = 0 corresponds to the animal pole, and *x* = ±*L* to the lateral sides. Our aim is to predict the thickness profile *h*(*x*) based on assuming a spatially patterned viscosity profile *η*(*x*).

To this end, we divide the fluid into *N* elements along the *x*-axis. Based on incompressibility, we assume that each fluid element *i* with thickness *hi* and length *ℓi* has a conserved area, and thus

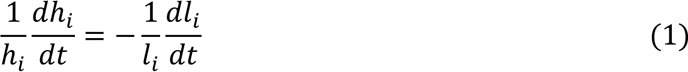

Assuming the constitutive equation for stress

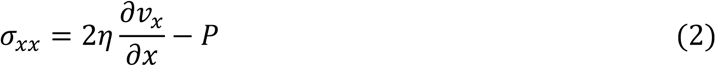

where *η* is the viscosity and *P* the pressure, we get

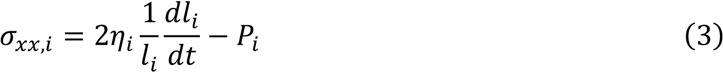

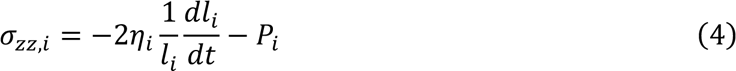

Applying force balance,

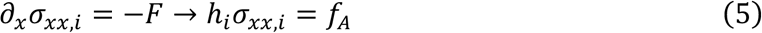

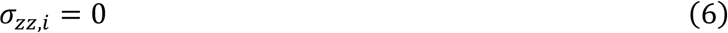

where *F* = *fA*/(*hiℓi*) is the force density. We then consider an active force that adjusts itself such that the extension speed is constant:

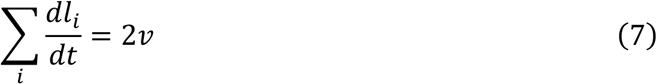

Putting everything together, we find

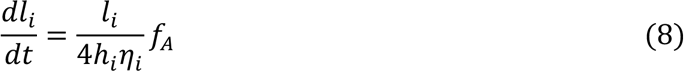

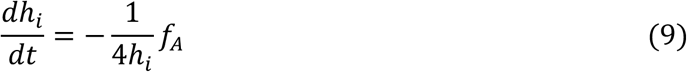

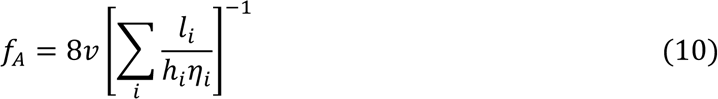

To compare model predictions to experiment, we use parameters corresponding to the experiment, *v* ≈ 1.5 μm/min^64^, *L* ≈ 500 µm. Finally, as a single free fit parameter, we estimate the ratio of maximal (lateral) viscosity *η*Lat = *η*(*x* = ±*L*) to minimal (animal) viscosity *η*An = *η*(*x* = 0) to be *η*Lat/*η*An ≈ 5.

## Acknowledgements

We are grateful to members of the Heisenberg lab for discussions, technical advice and feedback on the manuscript, to Irene Steccari and David Labrousse Arias for support in the generation of the plasmids used in this work, to Nicole Amberg and Florian Pauler for help and technical advice with the RNA sequencing, to Nicoletta Petridou, Elena Scarpa and Edouard Hannezo for critically reading the manuscript. We also thank the Imaging and Optics Facility, the Life Science facility and the Scientific Computing Unit at IST Austria for support. The RNA sequencing for animal and lateral ectoderm was performed by the Next Generation Sequencing Facility at Vienna BioCenter Core Facilities (VBCF), member of the Vienna BioCenter (VBC), Austria. D.B.B. was supported by the NOMIS Foundation as a NOMIS Fellow and by an EMBO Postdoctoral Fellowship (ALTF 343-2022). S.Tav. was supported by an EMBO Postdoctoral Fellowship (ALTF 1159-2018). A.S. is a recipient of a DOC Fellowship of the Austrian Academy of Science at IST Austria.

## Author contributions

Conceptualization, S.Tav. and C.-P.H; methodology, S.Tav. and D.B.B.; software S.Tas. and H.R.; formal analysis, S.Tav. and D.B.B.; investigation, S.Tav., D.B.B. and X.T.; resources, R.K.; writing – original draft, S.Tav. and C.-P.H.; writing – review & editing, S.Tav., D.B.B., S.Tas., X.T., A.S. and C.-P.H.; visualization, S.Tav.; funding acquisition, S.Tav., D.B.B., A.S. and C.-P.H.; project administration, C.-P.H.; supervision, C.-P.H.

---

**Supplementary Figure S1.**
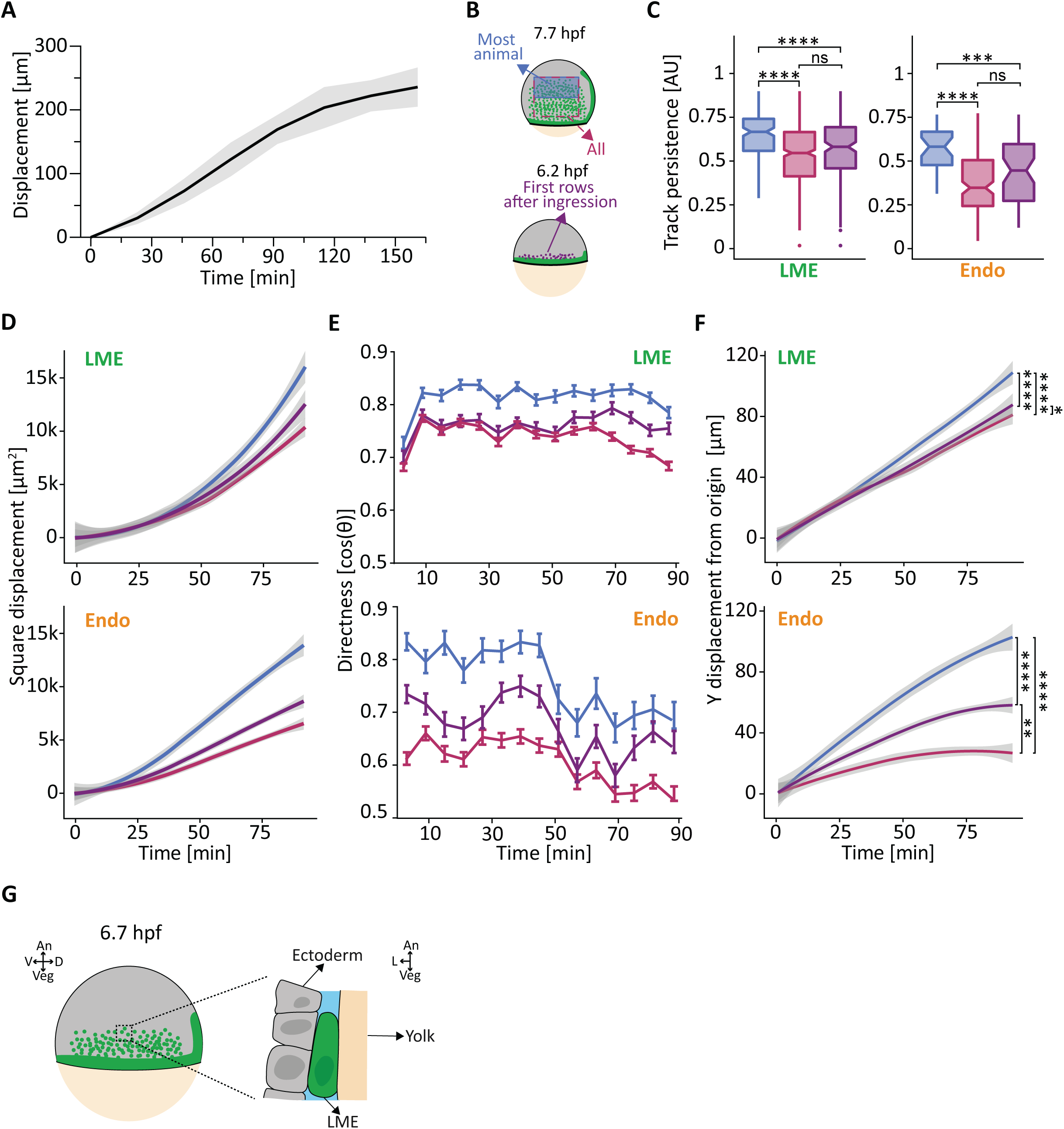
Lateral mesendoderm and lateral endoderm cells reaching the most animal part of the embryo undergo directed and persistent migration, related to Figure 1 (A) Displacement of the lateral mesendoderm (LME) front over time. The displacement of the LME front from the origin of migration towards the animal pole (Y-axis) was tracked every 23 min (t0 ∼6.2 hours post fertilization, hpf) for at least 115 min. Values are shown as mean (solid black line) with standard deviation (SD, light grey area). Number of embryos, 11. (B) Schematic representation of the three subpopulations of lateral mesendoderm (LME) and lateral endoderm (Endo) used for the migratory behaviour analyses in (C-F). Cells were tracked from 6.2 to 7.7 hpf (∼92 min total with ∼2 min frame rate). Upper part, cells were grouped in the subpopulation “All” (magenta; cells that were located within the 300 x 200 µm rectangular area outlined by magenta dashed lines at the end of tracking (92 min)) or “Most animal” (light blue; cells that were located within a 300 x 100 µm rectangular area marked by blue shadow at the end of tracking (92 min)) Lower part, cells were grouped in the subpopulation “First rows after ingression” (purple dots) when they were within the first 2-3 cell rows after ingression (start of tracking, 0 min). (C) Migration persistence of the “Most animal”, “All” and “First rows after ingression” subpopulations of LME (left box plot) and Endo (right box plot) cells. Number of LME cells: (Most animal, light blue), 150; (All, magenta), 446; (First rows after ingression, purple), 173. Number of embryos, 3. Number of Endo cells: (Most animal, light blue), 32; (All, magenta), 138; (First rows after ingression, purple), 54. Number of embryos, 2. Statistical test, Mann- Whitney test: ns not significant, ***p<0.001, ****p<0.0001. (D) Square Displacement of the “Most animal”, “All” and “First rows after ingression” subpopulations of LME (top plot) and Endo (bottom plot) cells, displayed as smooth plot. Solid line represents the mean, grey ribbon displays confidence interval. Number of LME cells: (Most animal, light blue), 150; (All, magenta), 446; (First rows after ingression, purple), 173. Number of embryos, 3. Number of Endo cells: (Most animal, light blue), 32; (All, magenta), 138; (First rows after ingression, purple), 54. Number of embryos, 2. (E) Directness of migration over time of the “Most animal”, “All” and “First rows after ingression” subpopulations of LME (top plot) and Endo (bottom plot) cells. The directness is calculated as angular deviation of the cell path [cos(8)] (where 1 equals to no deviation) from the predicted direct path (for more detailed information, see Methods). Values are shown as mean (solid line) with standard deviation (SD, error bars). Number of LME cells: (Most animal, light blue), 150; (All, magenta), 446; (First rows after ingression, purple), 173. Number of embryos, 3. Number of Endo cells: (Most animal, light blue), 32; (All, magenta), 138; (First rows after ingression, purple), 54. Number of embryos, 2. (F) Displacement towards the animal pole over time of the “Most animal”, “All” and “First rows after ingression” subpopulations of LME (top plot) and Endo (bottom plot) cells from. The displacement along the animal-vegetal axis is quantified as cumulative Y-displacement from the origin of migration (Y-axis) and displayed as smooth plot. Solid line represents the mean, grey ribbon displays confidence interval. Number of LME cells: (Most animal, light blue), 150; (All, magenta), 446; (First rows after ingression, purple), 173. Number of embryos, 3. Number of Endo cells: (Most animal, light blue), 32; (All, magenta), 138; (First rows after ingression, purple), 54. Number of embryos, 2. Statistical test on the final Y-displacement, Mann-Whitney test:, *p<0.05, **p<0.01, ****p<0.0001. (G) Schematic representation of an embryo during LME animal migration. The cross-section view (right) shows the interaction of LME with ectoderm and yolk cell. An, animal; Veg, vegetal; D, dorsal; V, ventral; L, lateral; hpf, hours post-fertilization.

**Supplementary Figure S2.**
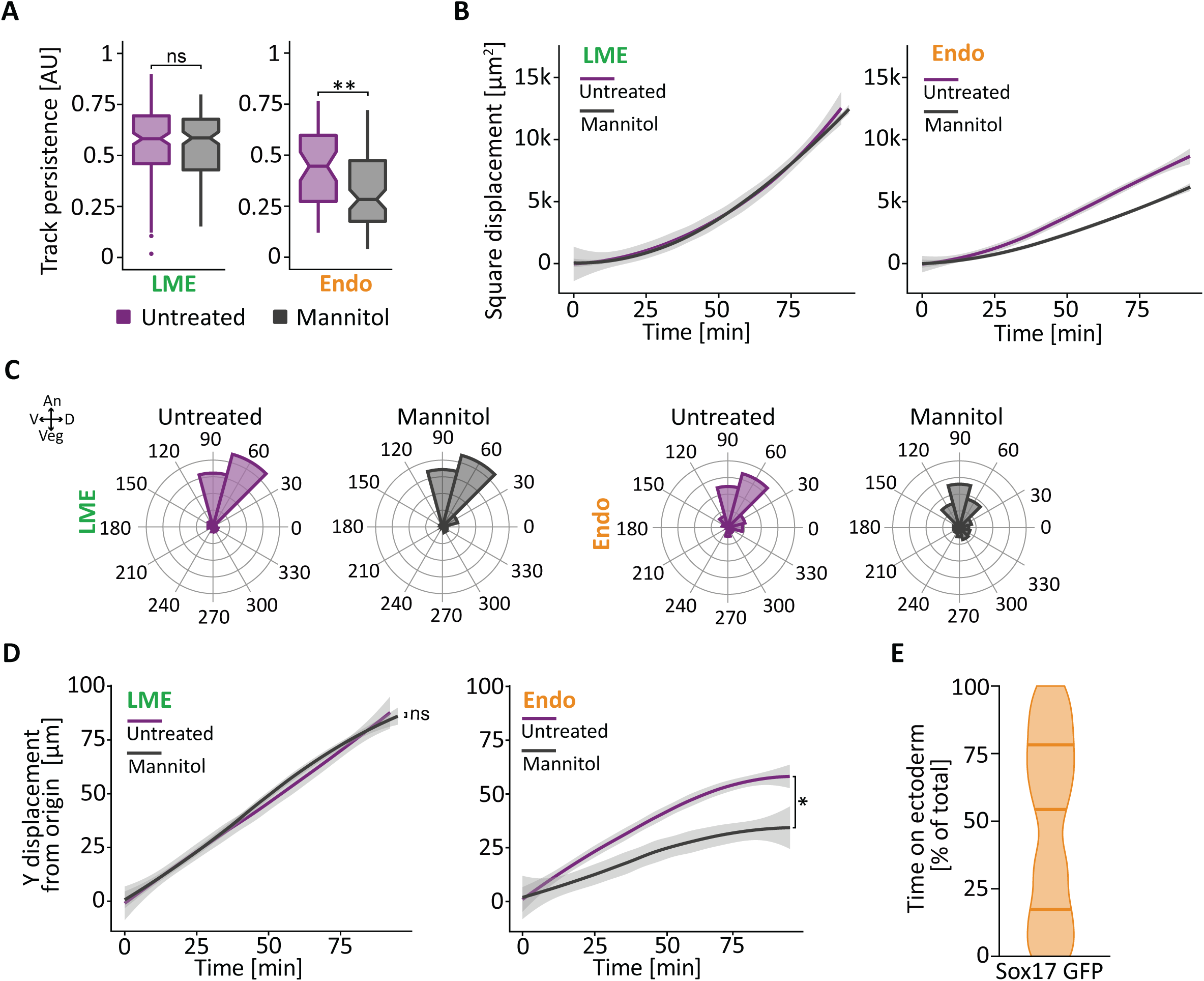
Mannitol-injection does not perturb lateral mesendoderm migration, related to Figure 1 (**A-E**) Sphere/dome stage (4-4.3 hours post fertilization, hpf) *Tg(sebox::eGFP);Tg(actb2::H2A- mCherry)* embryos (lateral mesendoderm (LME), A-D) or *Tg(sox17::eGFP)* (lateral endoderm (Endo) A-E) embryos injected with 50 pg *H2A-mCherry* mRNA at the 1-cell stage (to label nuclei), and 1 nl of 350 mM Mannitol (to increase interstitial fluid accumulation) together with 1.25 µg/µl 10.000 MW Alexa Fluor™ 647 dextran (to label interstitial fluid) were used. First ingressing cells (for definition see Figure S1B and Methods) were tracked for ∼92 min from the onset of animal pole-directed migration (∼2 min frame rate, t0 ∼6.2 hpf). (A) Migration persistence of the first ingressing lateral mesendoderm (LME) (left box plot) and lateral endoderm (Endo) (right box plot) cells in untreated control embryos and embryos injected with Mannitol. Number of LME cells: 173 from 3 untreated embryos (untreated, purple), 163 from 3 Mannitol-injected embryos (Mannitol, black). Number of Endo cells: 53 from 2 embryos (untreated, purple), 96 from 2 Mannitol-injected embryos (Mannitol, black). Dataset for untreated corresponds to data shown in Figure S1. Statistical test, Mann-Whitney test: ns not significant, ** p<0.01. (B) Square Displacement of the first ingressing LME (left box plot) and Endo (right box plot) cells in untreated control embryos and embryos injected with Mannitol, displayed as smooth plot. Solid line represents the mean, grey ribbon displays confidence interval. Number of LME cells: 173 from 3 untreated embryos (untreated, purple), 163 from 3 Mannitol-injected embryos (Mannitol, black); Number of Endo cells: 54 from 2 untreated embryos (untreated, purple), 96 from 2 Mannitol-injected embryos (Mannitol, black). Dataset for untreated corresponds to data shown in Figure S1. (C) Migration directionality of the first ingressing LME (left rose diagrams) and Endo (right rose diagrams) cells in untreated control embryos and embryos injected with Mannitol. Directionality is shown as rose diagrams where each concentric circle represents a different frequency range, with 0 at the centre and increasing by 10 for each subsequent circle. An, animal; Veg, vegetal; D, dorsal; V, ventral. Number of LME cells: 173 from 3 embryos (untreated, purple), 163 from untreated 3 Mannitol-injected embryos (Mannitol, black); Number of Endo cells: 54 from untreated 2 embryos (untreated, purple), 96 from 2 Mannitol- injected embryos (Mannitol, black). Dataset for untreated corresponds to data shown in Figure S1. (D) Displacement of the first ingressing LME (left plot) and Endo (right plot) cells in untreated control embryos and embryos injected with 350 mM Mannitol. The displacement along the animal-vegetal axis is quantified as cumulative Y-displacement from the origin of migration along the Y-axis and displayed as smooth plot. Solid line represents the mean, grey ribbon displays confidence interval. Number of LME cells: 173 from untreated 3 embryos (untreated, purple), 163 from 3 Mannitol-injected embryos (Mannitol, black). Number of Endo cells: 54 from untreated 2 embryos (untreated, purple), 95 from 2 Mannitol-injected embryos (Mannitol, black). Dataset for untreated corresponds to data shown in Figure S1. Statistical test on the final Y-displacement, Mann-Whitney test: ns not significant, * p<0.05. (E) Time Endo cells use the ectoderm as their substrate for migration. Endo cells were scored as either on the ectoderm or on the yolk membrane at each timepoint for ∼92 min from the onset of animal pole-directed migration. Data are shown as percentage of the total time of migration. Number of cells, 96. Number of embryos, 2.

**Supplementary Figure S3.**
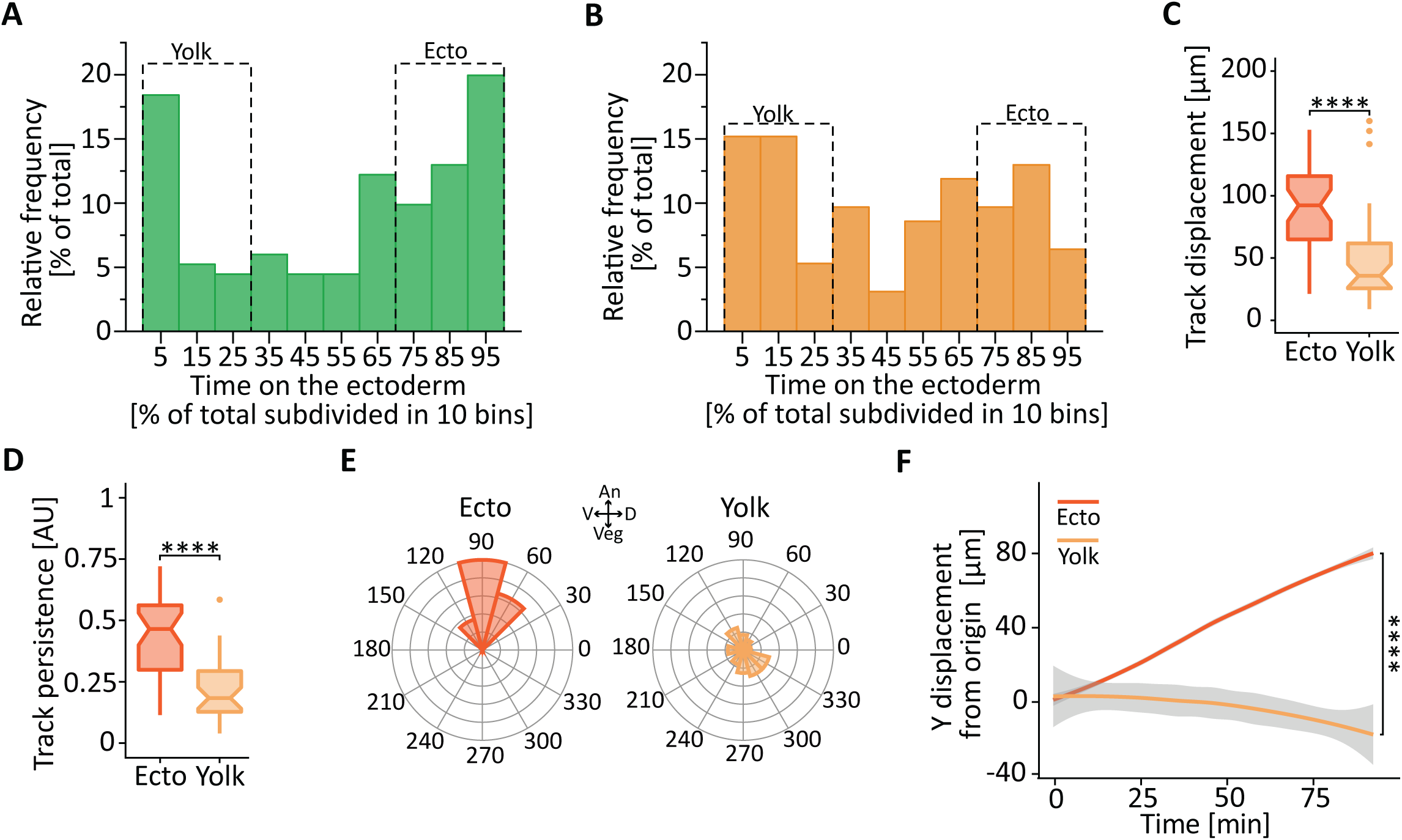
Lateral endoderm cells migrating on the ectoderm display most pronounced animal pole-directed migration, related to Figure 2 (**A-F**) Sphere/dome stage (4-4.3 hours post fertilization, hpf) *Tg(sebox::eGFP);Tg(actb2::H2A- mCherry)* embryos (lateral mesendoderm (LME), A) or *Tg(sox17::eGFP)* (lateral endoderm (Endo), B-F) embryos injected with 50 pg *H2A-mCherry* mRNA at the 1-cell stage to mark nuclei, and 1 nl of 350 mM Mannitol with 1.25 µg/µl 10.000 MW Alexa Fluor™ 647 dextran to label the interstitial fluid. First ingressing cells (for definition see Figure S1B and Methods) were tracked for ∼92 min from the onset of animal pole-directed migration (∼2 min frame rate, t0 ∼6.2 hpf). (**A,B**) Frequency distribution of LME (A) and Endo (B) cells as a function of time spent migrating on the ectoderm. The frequency (Y-axis) was calculated in bins of 10% of the total percentage of time spent on the ectoderm (X-axis). The dashed rectangles outline the cells using ectoderm as substrate for either more than 70% or less than 30% of their total migration time. These subgroups were subsequently used in the analysis as cells migrating mostly on the ectoderm (Ecto) or mostly on the yolk cell (Yolk), respectively. Number of LME cells: 163 from 3 embryos. Number of Endo cells: 96 from 2 embryos. (C) Total displacement of Endo cells using mostly the ectoderm (Ecto, dark orange) or the yolk cell (Yolk, light orange) as their substrate for migration. Number of cells: Ecto, 40; Yolk, 37. Number of embryos, 2. Statistical test, Mann-Whitney test: ****p<0.0001. (D) Migration persistence of Endo cells using mostly the ectoderm (Ecto, dark orange) or the yolk cell (Yolk, light orange) as their substrate for migration. Number of cells: Ecto, 40; Yolk, 37. Number of embryos, 2. Statistical test, Mann-Whitney test: ****p<0.0001. (E) Migration directionality of LME cells using mostly the ectoderm (Ecto, dark orange) or the yolk (Yolk, light orange) as their substrate for migration. Directionality is shown as rose diagram where each concentric circle represents a different frequency range, with 0 at the centre and increasing by 10 for each subsequent circle. An, animal; Veg, vegetal; D, dorsal; V, ventral. Number of cells: Ecto, 40; Yolk, 37. Number of embryos, 2. (F) Displacement of Endo cells towards the animal pole over time. The displacement along the animal-vegetal axis was quantified as cumulative Y-displacement from the origin of migration and displayed as smooth plot. Solid line represents the mean, grey ribbon displays confidence interval. Endo cells using mostly the ectoderm (Ecto) or the yolk cell (Yolk) as their substrate for migration are shown in dark and light orange, respectively. Number of cells: Ecto, 40; Yolk, 37. Number of embryos, 2. Statistical test on the final Y-displacement, Mann-Whitney test: ****p<0.0001.

**Supplementary Figure S4.**
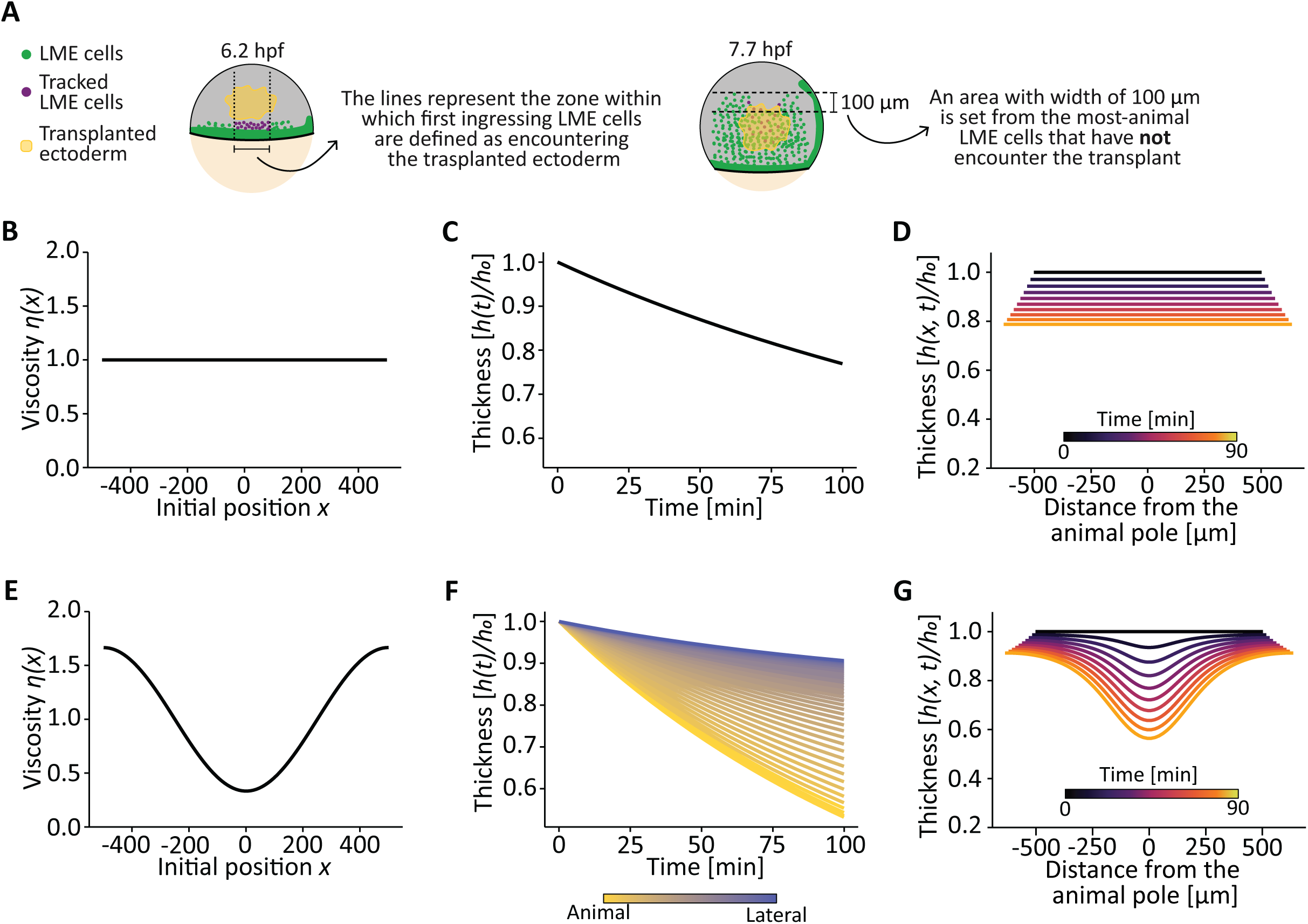
Minimal model of the ectoderm with patterned viscosity recapitulates the experimental ectodermal tissue thinning, related to Figures 3 and 4 (**A**) Schematic representation of the selection criteria for “first ingressing lateral mesendodermerm (LME) cells” (left) and definition of “most-animally migrating cells” (right) in transplanted embryos. On the left, to analyse the effect of the transplanted ectoderm (yellow shadowing) on LME migration, first ingressing LME cells (violet dots) were selected at the start of the tracking (6.2 hours post fertilization, hpf) within an area (black dotted lines), which was vegetal to the transplanted ectoderm and 1-2 cell diameters smaller on both sides in comparison to the transplant itself. On the right, to quantify the most-animally migrating cells at the end of the tracking (7.7 hpf), the most-animally-located LME cells that have not encountered the transplanted ectoderm (yellow shadowing) were used to set the animal boundary of 100 μm-wide bin (dashed lines). Tracked LME cells (violet dots) that were found within the bin were scored as most-animally migrating LME cells. (**B-G**) Model of the ectoderm thickness as a function of developmental time. Ectoderm tissue was modelled as a passive fluid with either constant viscosity (B-D) or patterned viscosity from the animal pole (lowest) to the margin (highest) (E-F) and subjected to an external constant pulling force, which cause an increase of tissue length with a speed of ∼1 µm/min (corresponding to the experimentally measured epiboly speed). Ectoderm thickness in (C, D, F and G) was quantified as function of time. In (D and G), the x-axis represents the position relative to the animal pole. The initial length (t0) of the ectoderm tissue was set at 1000 µm.

**Supplementary Figure S5.**
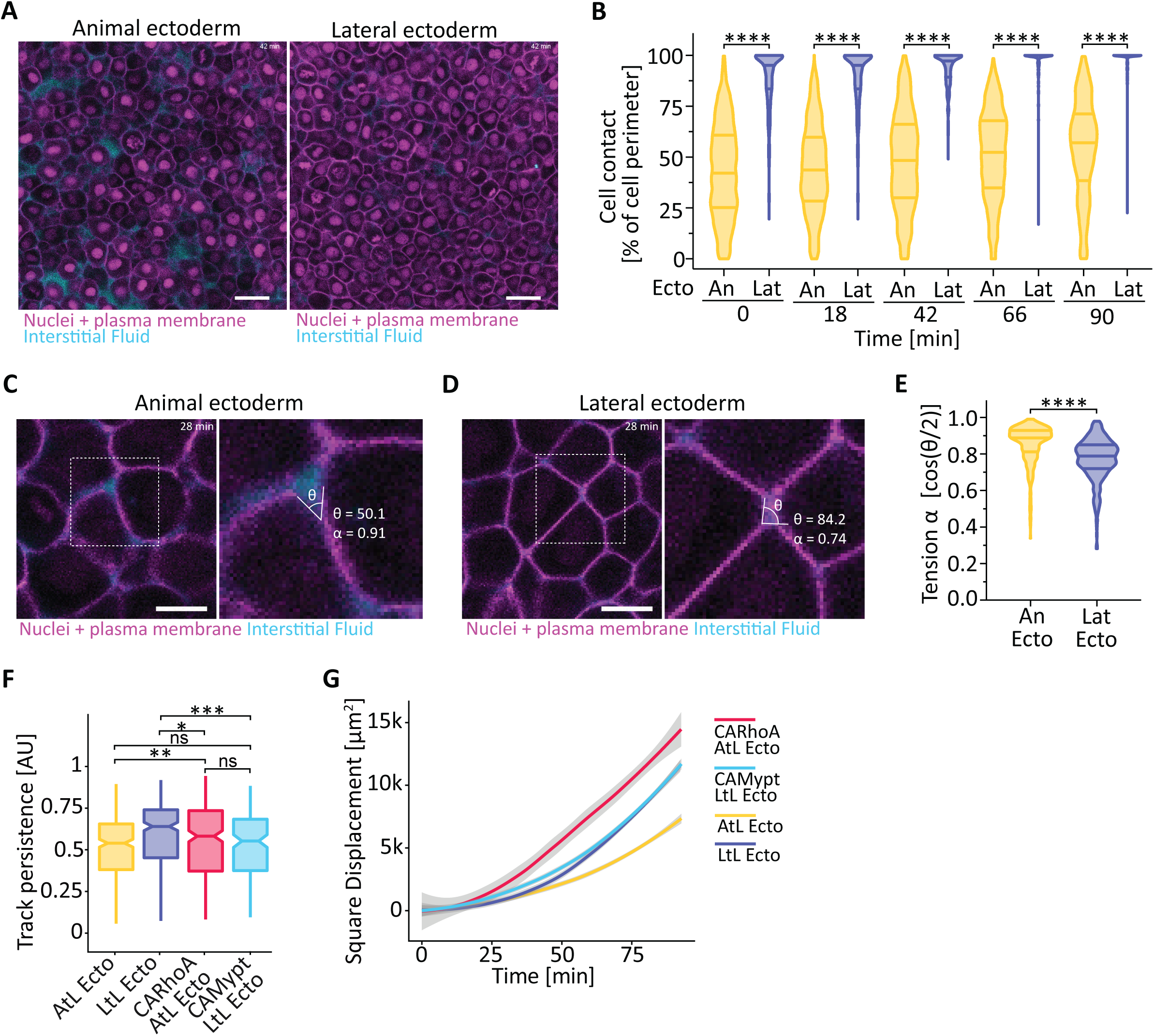
Lateral ectoderm cells display a higher percentage of cell perimeter engaged in cell-cell contacts, related to Figures 5 and 6 (A) Fluorescence images of animal (left) and lateral (right) ectoderm from wildtype *Tg(sebox::eGFP)* embryos injected with 35 pg *H2A-mCherry* (nuclei) and 100 pg *membrane RFP* (cell membrane) mRNAs at the 1-cell stage. At sphere/dome stage (4-4.3 hours post fertilization, hpf), embryos were injected with 1nl of 1.5 µg/µl 10.000 MW Cascade Blue™ dextran to mark interstitial fluid. Images are maximum intensity projection of stacks of three 1 µm optical sections (Z step size, 1.5 µm). Magenta, H2A-mCherry and membrane RFP (nuclei and cell membrane); cyan, Cascade Blue™ dextran (interstitial fluid). Scale bar, 25 μm. See also Video S10. (B) Cell-cell contacts in the animal and lateral ectoderm as a function of time with 0 min ∼6.2 hpf. Cell contacts are depicted as percentage of the total cell perimeter (Y-axis). Number of cells: animal ectoderm (An, yellow), 528-615 from 4 embryos; lateral ectoderm (Lat, blue), 271-397 from 2 embryos. Statistical test, Mann-Whitney test: ****p<0.0001. (**C,D**) Constitutive active RhoA (CARhoA) overexpressing animal-to-lateral and constitutive active Mypt (CAMypt) overexpressing lateral-to-lateral ectoderm transplant assay. Animal ectoderm cells (red, A) and lateral ectoderm cells (light blue, B) from a *Tg(sebox::eGFP)* donor embryo injected with either 1-2 pg *CARhoA* mRNA (A) or 75 pg *CAMypt* mRNA (B) together with 10-15 pg *H2B-mKO2k* and 100 pg *membrane RFP* mRNAs to mark nuclei and plasma membrane, respectively, were transplanted at germ ring stage (5.7 hpf) into the lateral ectoderm of a stage-matched wildtype *Tg(sebox::eGFP)* host embryo injected with 10-15 pg *H2B-mKO2k* mRNA (nuclei). After transplantation, lateral mesendoderm (LME) cells encountering the transplanted ectoderm were tracked for ∼92 min (∼2 min 14 sec frame rate, t0 ∼6.2 hpf). (**C,D**) Fluorescence images of animal (C) and lateral (D) ectoderm from wildtype *Tg(sebox::eGFP); Tg(actb2::H2B-mCherry)* embryos injected 100 pg *membrane RFP* (cell membrane) mRNAs at the 1-cell stage. At sphere/dome stage (4-4.3 hours post fertilization, hpf), embryos were injected with 1nl of 1 µg/µl 10.000 MW Alexa Fluor™ 647 to mark interstitial fluid. Dashed area demarcates the area shown in the left panels with exemplary contact angles (8) used to determine the relative cell tension (α). The images are single 1 µm-thick optical sections. Magenta, H2B-mCherry and membrane RFP (nuclei and cell membrane); cyan, Alexa Fluor™ 647 dextran (interstitial fluid). Scale bar, 10 μm. (**D**) Relative cell tension in the animal and lateral ectoderm 28 min after the onset of LME migration (∼6.2 hpf). The relative cell tension is calculate as cosine of the contact angle in radiant divided by two. Number of contact angles: animal ectoderm (An Ecto, yellow), 343 from 2 embryos; lateral ectoderm (Lat Ecto, blue), 378 from 2 embryos. Statistical test, Mann- Whitney test: ****p<0.0001. (**F**) Migration persistence of LME cells in transplanted embryos. Number of cells: 292 from 6 CARhoA overexpressing animal-to-lateral ectoderm transplants (CARhoA AtL Ecto, red); 162 from 4 CAMypt overexpressing lateral-to-lateral ectoderm transplants (CAMypt LtL Ecto, light blue); 327 from 9 animal-to-lateral ectoderm transplants (AtL Ecto, yellow); 327 from 9 lateral-to-lateral ectoderm transplants (LtL Ecto, blue). Datasets for AtL Ecto and LtL Ecto correspond to data shown in Figure 3. Statistical test, Mann-Whitney test: ns not significant, * p<0.05, ** p<0.01, *** p<0.001. (**G**) Square Displacement of LME cells in transplanted embryos, displayed as smooth plot. Solid line represents the mean, grey ribbon displays confidence interval. Number of cells: 292 from 6 CARhoA overexpressing animal-to-lateral ectoderm transplants (CARhoA AtL Ecto, red); 162 from 4 CAMypt overexpressing lateral-to-lateral ectoderm transplants (CAMypt LtL Ecto, light blue); 327 from 9 animal-to-lateral ectoderm transplants (AtL Ecto, yellow); 327 from 9 lateral-to-lateral ectoderm transplants (LtL Ecto, blue). Datasets for AtL Ecto and LtL Ecto correspond to data shown in Figure 3.

**Supplementary Figure S6.**
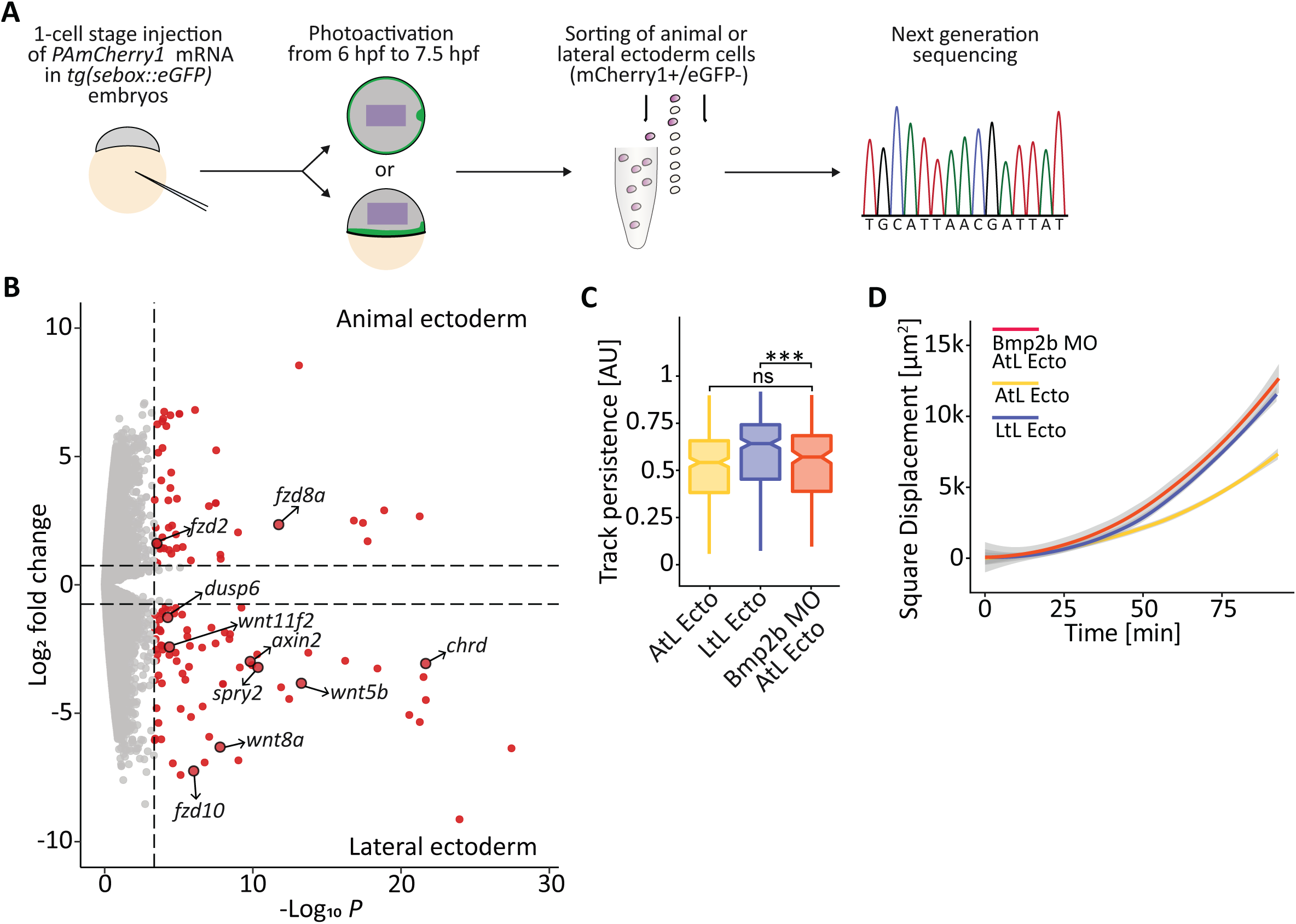
Transcriptomic analysis of lateral and animal ectoderm during lateral mesendoderm animal migration, related to Figure 7 (A) Schematic representation of bulk RNA sequencing (RNAseq) of the lateral and animal ectoderm in *Tg(sebox::eGFP)* embryos injected with 125 pg of *PAmCherry1* at the one-cell stage. For marking animal or lateral ectoderm, PAmCherry1 in the respective areas was photoactivated using a 405 nm laser from 6-7.5 hours post fertilization (hpf), followed by cell sorting, total mRNA extraction and sequencing. Number of samples: animal ectoderm,4; lateral ectoderm, 4. Number of embryos per sample: 28-32. (B) Differentially-expressed genes in animal and lateral ectoderm at 7.5 hpf. Volcano plot of all genes expressed in animal and lateral ectoderm with differentially-expressed genes marked in red. Genes more highly expressed in the animal ectoderm are in the upper part of the plot (positive log2 fold change), whereas genes more highly expressed in the lateral ectoderm are in lower part of the plot (negative log2 fold change). Differentially-expressed genes of signalling pathway components (except transcriptional regulators) are indicated with black arrows. See also Table S1. (**C,D**) *Bmp2b* morphant animal-to-lateral ectoderm transplant assay. Animal ectoderm cells (orange) from a *Tg(sebox::eGFP)* donor embryo injected with 1.5 ng *bmp2b morpholino* (MO) together with 15 pg *H2B-mKO2k* and 100 pg *membrane RFP* mRNAs to mark nuclei and plasma membrane, respectively, were transplanted at germ ring stage (5.7 hpf) into the lateral ectoderm of a stage-matched wildtype *Tg(sebox::eGFP)* host embryo injected with 10-15 pg *H2B-mKO2k* mRNA (nuclei). After transplantation, lateral mesendoderm (LME) cells encountering the transplanted ectoderm (for definition, see Figure S4 and Methods) were tracked for ∼92 min (∼2 min 14 sec frame rate, t0 ∼6.2 hpf). (C) Migration persistence of LME cells in transplanted embryos. Number of cells: 199 from 5 *bmp2b* MO animal-to-lateral ectoderm transplants (Bmp2b MO AtL Ecto, orange); 327 from 9 animal-to-lateral ectoderm transplants (AtL Ecto, yellow); 327 from 9 lateral-to-lateral ectoderm transplants (LtL Ecto, blue). Datasetss for AtL Ecto and LtL Ecto correspond to data shown in Figure 3. Statistical test, Mann-Whitney test: ns not significant, *** p<0.001. (D) Square Displacement of LME cells in transplanted embryos, displayed as smooth plot. Solid line represents the mean, grey ribbon displays confidence interval. Number of cells: 199 from 5 *bmp2b* MO animal-to-lateral ectoderm transplants (Bmp2b MO AtL Ecto, orange); 327 from 9 animal-to-lateral ectoderm transplants (AtL Ecto, yellow); 327 from 9 lateral-to-lateral ectoderm transplants (LtL Ecto, blue). Datasets for AtL Ecto and LtL Ecto correspond to data shown in Figure 3.

Table S1. Differentially-expressed genes in animal and lateral ectoderm at 7.5 hpf.

List of all differentially-expressed genes between animal and lateral ectoderm. Genes encoding for signalling pathway components (except transcriptional regulators) are in bold.

Video S1. Animal migration and tumbling phases of lateral mesendoderm cells, related to Figure 1.

Lateral mesendoderm (LME) migration in a *Tg(sebox::eGFP);Tg(actb2::mKate2-tpm3)* embryo from the onset of animal pole-directed migration (6.2 hours post fertilisation (hpf)) until the end of tumbling phase (8.8 hpf). LME, green; actin cytoskeleton, grey. Dashed lines in lateral view of the embryo (left) indicate the 20 μm-wide area used to create the cross-section view (right). ∼5 min 46 sec frame rate. Scale bar, 100 μm.

Video S2. Lateral mesendoderm cells sparsely labelled with Lifeact-RFP undergoing animal migration, related to Figure 1.

Lateral mesendoderm (LME) animal migration (6.2-7.2 hours post fertilisation (hpf)) in a *Tg(sebox::eGFP)* embryo sparsely labelled by injection of 7.5 pg *LifeAct-RFP* mRNA into a single blastomere at the 64/128 cells stage (2-2.25 hpf) to quantify protrusion directionality during animal migration. LME, green; actin cytoskeleton, grey; interstitial fluid, cyan. Arrows point to exemplary cells used for protrusion directionality analysis. Rectangular area outlined by white dashed lines in the yz view (left) demarcates the volume shown in the middle xy view. ∼2 min 14 sec frame rate. Scale bar, 20 μm.

Video S3. Lateral mesendoderm cells undergoing animal migration after injection of 350 mM Mannitol, related to Figure 1.

Lateral mesendoderm (LME) undergoing animal migration (6.2-7.5 hours post fertilisation (hpf)) in a *Tg(sebox::eGFP);Tg(actb2::H2A-mCherry)* embryo after injection at sphere/dome stage (4-4.3 hpf) of 1 nl of 350 mM Mannitol together (to increase interstitial fluid accumulation) with 1.25 µg/µl 10.000 MW Alexa Fluor™ 647 dextran (to mark the interstitial fluid). LME, green; nuclei, magenta; interstitial fluid, cyan. Dashed lines in the lateral view of the embryo indicate the position of the single plane yz (right) and xz (bottom) cross-section views. 2 min frame rate. Scale bar, 100 μm.

Video S4. Animal-to-lateral ectoderm transplant, related to Figure 3.

Lateral mesendoderm (LME) animal migration (6.2-7.5 hours post fertilisation (hpf)) in a *Tg(sebox::eGFP)* embryo after animal-to-lateral (heterotypic) ectoderm transplantation. LME, green; transplanted animal ectoderm, yellow. Dashed line in the lateral view of the embryo indicates the position of the single plane yz cross-section view. ∼2 min 20 sec frame rate. Scale bar, 100 μm.

Video S5. Lateral-to-lateral ectoderm transplant, related to Figure 3.

Lateral mesendoderm (LME) animal migration (6.2-7.5 hours post fertilisation (hpf)) in a *Tg(sebox::eGFP)* embryo after lateral-to-lateral (homotypic) ectoderm transplantation. LME, green; transplanted lateral ectoderm, blue. Dashed line in the lateral view of the embryo indicates the position of the single plane yz cross-section view. ∼2 min 20 sec frame rate. Scale bar, 100 μm

Video S6. Ectoderm thinning, animal pole view, related to Figure 4.

Thinning of the ectoderm during LME animal migration. Top panels, animal pole cross-section view of a *Tg(sebox::eGFP);Tg(actb2::mKate2-tpm3)* embryo during lateral mesendoderm animal migration (6.2-7.5 hours post fertilisation (hpf)). LME, green; actin cytoskeleton, magenta; interstitial fluid, cyan. Bottom panels, binary masks of the embryo (magenta), interstitial fluid (IF, cyan) and LME (green), generated from the cross-section view to quantify the thickness of the ectoderm during LME animal migration phase. ∼8 min 40 sec frame rate. Scale bar, 100 μm.

Video S7. Ectoderm thinning, lateral side view, related to Figure 4.

Thinning of the ectoderm during LME animal migration. Left panels, lateral side cross-section view of a *Tg(sebox::eGFP);Tg(actb2::mKate2-tpm3)* embryo during lateral mesendoderm animal migration (6.2-7.5 hours post fertilisation (hpf)). LME, green; actin cytoskeleton, magenta; interstitial fluid, cyan. Right, binary masks of the embryo (magenta), interstitial fluid (IF, cyan) and LME (green), generated from the cross-section view to quantify the thickness of the ectoderm during LME animal migration phase. ∼8 min 40 sec frame rate. Scale bar, 100 μm.

Video S8. Cell fraction in the animal and lateral ectoderm, related to Figure 5.

Cross-section views (xy (left) and yz (right)) of the animal (upper panels) and lateral (lower panels) ectoderm of *Tg(sebox::eGFP)* embryos injected with 35 pg *H2A-mCherry* mRNA (to mark nuclei) at the 1-cell stage and with 1nl of 1 µg/µl 10.000 MW Alexa Fluor™ 647 dextran (to mark interstitial fluid) at sphere/dome stage (4-4.3 hours post fertilization, hpf) used to quantify the cell fraction during lateral mesendoderm (LME) animal migration. LME, green; nuclei, magenta; interstitial fluid, cyan. Dashed line in the xy view (left) indicates the position of the yz cross-section view. Scale bar 20 μm.

Video S9. Cell fraction in the animal-to-lateral and lateral-to-lateral ectoderm transplants, related to Figure 5.

Cross-section views of the animal-to-lateral (left panel) and lateral-to-lateral (right panel) transplants in *Tg(sebox::eGFP)* host embryos injected with 1nl of 1 µg/µl 10.000 MW Alexa Fluor™ 647 dextran (tp marl interstitial fluid) at germ ring stage (5.7 hpf) during lateral mesendoderm (LME) animal migration. Transplanted ectoderm is from from *Tg(sebox::eGFP)* donor embryos injected with 35 pg *H2A-mCherry* mRNA (to mark nuclei) at the 1-cell stage. LME, green; transplanted animal ectoderm nuclei, yellow; transplanted lateral ectoderm nuclei, blue; interstitial fluid, cyan. Scale bar 20 μm.

Video S10. Cell-cell contacts in the animal and lateral ectoderm, related to Figure S5.

Cross-section views of the animal (left panels) and lateral (right panels) ectoderm of *Tg(sebox::eGFP)* embryos injected with 35 pg *H2A-mCherry* mRNA (to mark nuclei) and 100 pg *membrane RFP* (to mark cell membrane) mRNAs at the 1-cell stage and with 1nl of 1.5 µg/µl 10.000 MW Cascade Blue™ dextran (to mark interstitial fluid) at sphere/dome stage (4- 4.3 hours post fertilization, hpf) used for quantifying the percentage of cell perimeter engaged in cell-cell contact during lateral mesendoderm animal migration. Nuclei and cell membrane, magenta; interstitial fluid, cyan. Scale bar 20 μm

Video S11. CARhoA animal-to-lateral ectoderm transplant, related to Figure 6.

Lateral mesendoderm (LME) animal migration (6.2-7.5 hours post fertilisation (hpf)) in a *Tg(sebox::eGFP)* embryo after animal-to-lateral (heterotypic) transplantation of constitutive active RhoA (CARhoA) overexpressing ectoderm. LME, green; transplanted CARhoA animal ectoderm, red. Dashed line in the lateral view of the embryo indicates the position of the single plane yz cross-section view. ∼2 min 20 sec frame rate. Scale bar, 100 μm.

Video S12. CAMypt lateral-to-lateral ectoderm transplant. Transplanted lateral ectoderm is in light blue, LME in green. Related to Figure 6.

Lateral mesendoderm (LME) animal migration (6.2-7.5 hours post fertilisation (hpf)) in a *Tg(sebox::eGFP)* embryo after lateral-to-lateral (homotypic) transplantation of constitutive active Mypt (CAMypt) overexpressing ectoderm. LME, green; transplanted lateral ectoderm, light blue. Dashed line in the lateral view of the embryo indicates the position of the single plane yz cross-section view. ∼2 min 20 sec frame rate. Scale bar, 100 μm

Video S13. Lateral mesendoderm migration in control and Bmp2b-depleted embryos, related to Figure 7.

Animal pole view of lateral mesendoderm migration from 5.5 to 9.5 hours post fertilization (hpf) in *Tg(sebox::eGFP)* embryos injected with 1.5 ng control (Ctrl) (upper part) or 1.5 ng *bmp2b morpholino* (Bmp2b-depleted) (lower part). Mesendoderm, green; bright field, grey. 2 min frame rate. Scale bar, 100 μm.

**Video S14. Bmp2b morphant animal-to-lateral ectoderm transplant, related to Figure 7**. Lateral mesendoderm (LME) animal migration (6.2-7.5 hours post fertilisation (hpf)) in a *Tg(sebox::eGFP)* embryo after animal-to-lateral (heterotypic) transplantation of *Bmp2b* morphant ectoderm. LME, green; transplanted *Bmp2b* morphant animal ectoderm, orange. Dashed line in the lateral view of the embryo indicates the position of the single plane yz cross-section view. ∼2 min 20 sec frame rate. Scale bar, 100 μm.

